# Proteins with prion-like domains can form viscoelastic condensates that enable membrane remodeling and endocytosis

**DOI:** 10.1101/145664

**Authors:** Louis-Philippe Bergeron-Sandoval, Sandeep Kumar, Hossein Khadivi Heris, Catherine Chang, Caitlin E. Cornell, Sarah L. Keller, Paul François, Adam G. Hendricks, Allen J. Ehrlicher, Rohit V. Pappu, Stephen W. Michnick

## Abstract

Membrane invagination and vesicle formation are key steps in endocytosis and cellular trafficking. Here, we show that endocytic coat proteins with prion-like domains (PLDs) form hemispherical puncta in the budding yeast, *S. cerevisiae*. These puncta have the hallmarks of biomolecular condensates and enable membrane remodeling to drive actin-independent endocytosis. The puncta, which we refer to as endocytic condensates, form and dissolve reversibly in response to changes in temperature and solution conditions. The condensates are organized around dynamic protein-protein interaction networks, which involve interactions among PLDs with high glutamine contents. The endocytic coat protein Sla1 is at the hub of the protein-protein interaction network. Using active rheology, we indirectly characterized the material properties of endocytic condensates. These experiments show that endocytic condensates are viscoelastic materials and allow us to estimate the interfacial tension between endocytic condensates and their surroundings. We then adapt the physics of contact mechanics, specifically the contact theory of Hertz, to develop a quantitative framework for describing how interfacial tensions among condensates, the membrane, and the cytosol can deform the plasma membrane to enable actin independent endocytosis.

## Introduction

Endocytosis in eukaryotic cells can occur via two separate mechanisms. In the budding yeast *Saccharomyces cerevisiae*, membrane invagination that enables endocytosis is normally driven by growth of membrane-bound branched actin (Carlsson and Bayly 2014). A second actin independent route to endocytosis is realized when intracellular turgor pressure is reduced. This reduction of turgor pressure alleviates the tension on plasma membranes that would normally oppose membrane invagination (Aghamohammadzadeh and Ayscough 2009, Basu, Munteanu et al. 2014). In both mechanisms, endocytosis is initiated by the coordinated recruitment of a number of proteins associated with distinct stages of endocytic maturation (Kaksonen, Toret et al. 2005). Clathrin heavy and light chains first interact with initiator proteins (Ede1 and Syp1) to form a lattice on the membrane. Subsequently, early coat proteins such as Sla1, Sla2, Ent1, Ent2, and Yap1801 (Malinovska, Kroschwald et al. 2013) bind directly to the adaptor-clathrin lattice and form the cortical body (Kaksonen, Toret et al. 2005).

Electron microscopy data highlight the existence of hemispherical membraneless bodies around endocytic sites. These bodies are identifiable by following the localization of labeled endocytic coat proteins such as Sla1. The observed Sla1-labeled bodies are known to exclude ribosomes from the cytosol near the cortical sites. Importantly, these endocytic bodies form even when actin is not polymerized and the membrane is flat (Kukulski, Schorb et al. 2012). Many of the coat proteins in bodies that form around endocytic sites include prion-like domains (PLDs). These domains are intrinsically disordered, low complexity sequences that are enriched in polar amino acids such as glutamine, asparagine, glycine, and serine and are interspersed by aromatic residues such as tyrosine (Alberti, Halfmann et al. 2009, Malinovska, Kroschwald et al. 2013).

Proteins with PLDs have the ability to undergo phase separation in cells (Alberti, Saha et al. 2018) and *in vitro* (Martin, Holehouse et al. 2020) giving rise to the formation of biomolecular condensates that are mesoscale, non-stoichiometric macromolecular assemblies that concentrate biomolecules (Bergeron-Sandoval, Safaee et al. 2016, Banani, Lee et al. 2017, Powers, Holehouse et al. 2019). Here, we show that endocytosis in *S. cerevisiae* involves the concentration of PLD-containing proteins – such as the essential protein Sla1 – within biomolecular condensates that form at cortical sites. These condensates are viscoelastic, and they are scaffolded by a dense network of PLD-containing proteins. We find that condensate formation requires an intact PLD and that the coat protein Sla1 is at the hub of the condensate driving protein-protein interaction network. The distinctive compositional biases within PLDs of coat proteins contribute to condensate formation and function. We show that the formation of the condensates and cohesiveness of molecular interactions within them, are essential for mechanoactive processes associated with actin-independent endocytosis. We present a model, motivated by the contact theory of Hertz (Hertz 1881, Derjaguin, Muller et al. 1975, Popov, Pohrt et al. 2017), to provide a plausible explanation for how interfacial tensions among condensates, the membrane, and the cytosol, can enable membrane invagination and drive actin-independent endocytosis.

## Results

### Sla1-labeled puncta are characterized by temperature-dependent non-covalent interactions

We sought to determine the role of PLD-containing early coat proteins in endocytosis. In the budding yeast *S. cerevisiae* we investigated actin-independent endocytosis by deleting the gene that codes for glycerol-3-phosphate dehydrogenase, a key enzyme of glycerol synthesis, (*GPD1Δ*), (Basu, Munteanu et al. 2014). Glycerol is the major osmolyte in yeast cells, and glycerol deficiency alleviates turgor pressure and tension on the membrane in *GPD1Δ* cells (*SI Appendix*, Fig. S1*A*). We could then interrogate actin-independent endocytosis in *GPD1Δ* cells by treating them with Latrunculin A (Lat A), an inhibitor of actin polymerization. The concentrations of Lat A that we use in our experiments cause a loss in localization of the filamentous actin marker Abp1 (*SI Appendix*, Fig. S1 *B-C*) (Aghamohammadzadeh, Smaczynska-de et al. 2014). We then tracked the formation and internalization of Sla1-labeled endocytic puncta at cortical sites (*SI Appendix*, Fig. S1*D*).

Proteins with PLDs and many PLDs themselves undergo reversible, thermoresponsive phase transitions with upper critical solution temperatures (UCST) in cells (Alberti, Saha et al. 2018) and *in vitro* (Martin, Holehouse et al. 2020). Systems characterized by UCST phase behavior will undergo phase separation below a critical temperature Tc and transition to one-phase homogeneous regimes above Tc (Ruff, Roberts et al. 2018). The ability of a biomolecular condensate to undergo an abrupt and reversible switch between two phases at a distinct temperature, and do so over multiple heating and cooling cycles, is a defining feature of reversible, thermoresponsive phase transitions (Rayermann, Rayermann et al. 2017). Motivated by published results for PLDs and proteins with PLDs, we assessed whether the Sla1-labeled endocytic puncta also form and dissolve reversibly in response to increases / decreases to temperature. To investigate such behavior for endocytic puncta, we quantified their sensitivity to multiple cycles of heating and cooling. Sla1-labeled puncta in *GPD1Δ* cells dissolved and reassembled within seconds as the temperature of the growth medium was cycled between 30°C and 49°C (Fig. 1*A-C*). Quantification of the number of puncta per cell showed a bistable response to increases / decreases in temperature with an apparent transition temperature (T_t_) of 45.3°C. Specifically, cells did not contain puncta above T_t_ whereas puncta were observed in cells below T_t_ (Fig. 1*A, B*). Importantly, cells remained viable above T_t_ and endocytic function was restored when the temperature dropped below T_t_ (*SI Appendix*, Fig. S2*A, B*).

**Figure 1.**
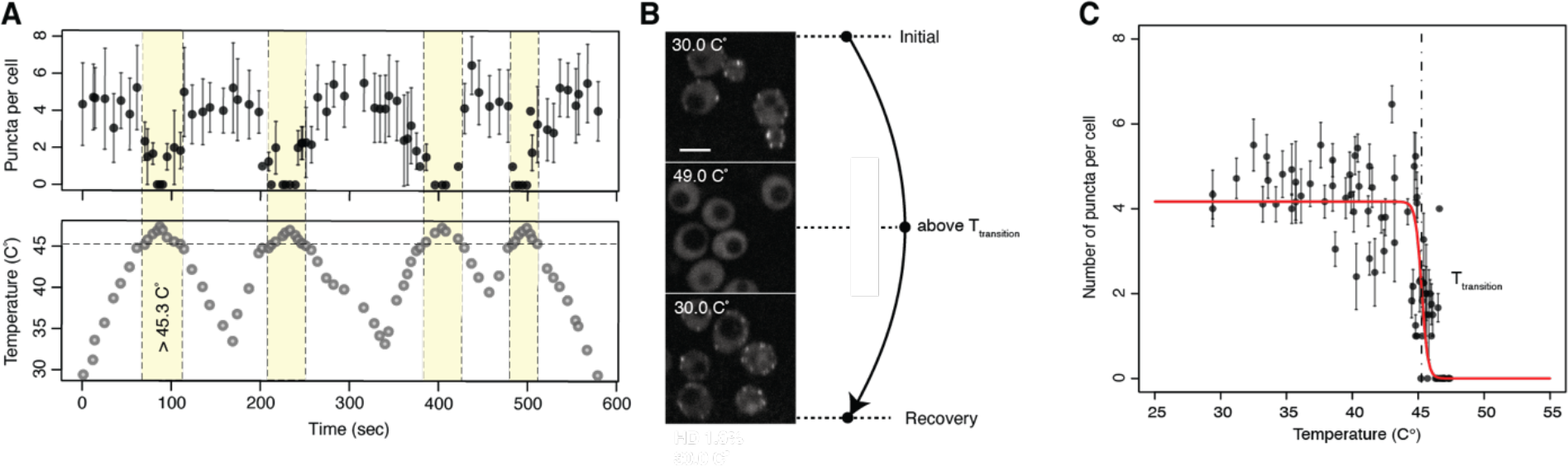
Endocytic proteins with PLDs form thermo-sensitive puncta. (A) Reversible, temperature-dependent dissolution and formation of Sla1-labeled endocytic puncta when cells are cycled above and below an apparent transition temperature (T_t_) of 45.3 C° (yellow region) over multiple heating and cooling cycles (left; mean ± sd; n > 8 cells). (B) Representative images of temperature-dependent dissolution and reformation of Sla1-labeled endocytic puncta in *GPD1Δ* cells treated with Lat A when cells are cycled above and below a transition temperature (T_t_) over a heating and cooling cycle. Scale bar, 4 μm. (C) Average number of puncta per cell is bistable as a function of temperature.

### Proteins are labile within endocytic condensates

Using super-resolution fluorescence imaging, we quantified the dimensions of Sla1-labeled bodies to be 209 ± 10 nm long and 118 ± 6 nm wide. These inferences were based on the orientation of puncta in the imaging plane (Fig. 2*A*). Our observations for top projections are similar to previous reports of Sla-labeled spherical structures (Mund, van der Beek et al. 2018). They are also consistent with the electron microscopy data highlighting hemispherical bodies that exclude ribosomes from the cytosol near cortical sites that form in a manner that is independent of actin polymerization (Kukulski, Schorb et al. 2012).

**Figure 2.**
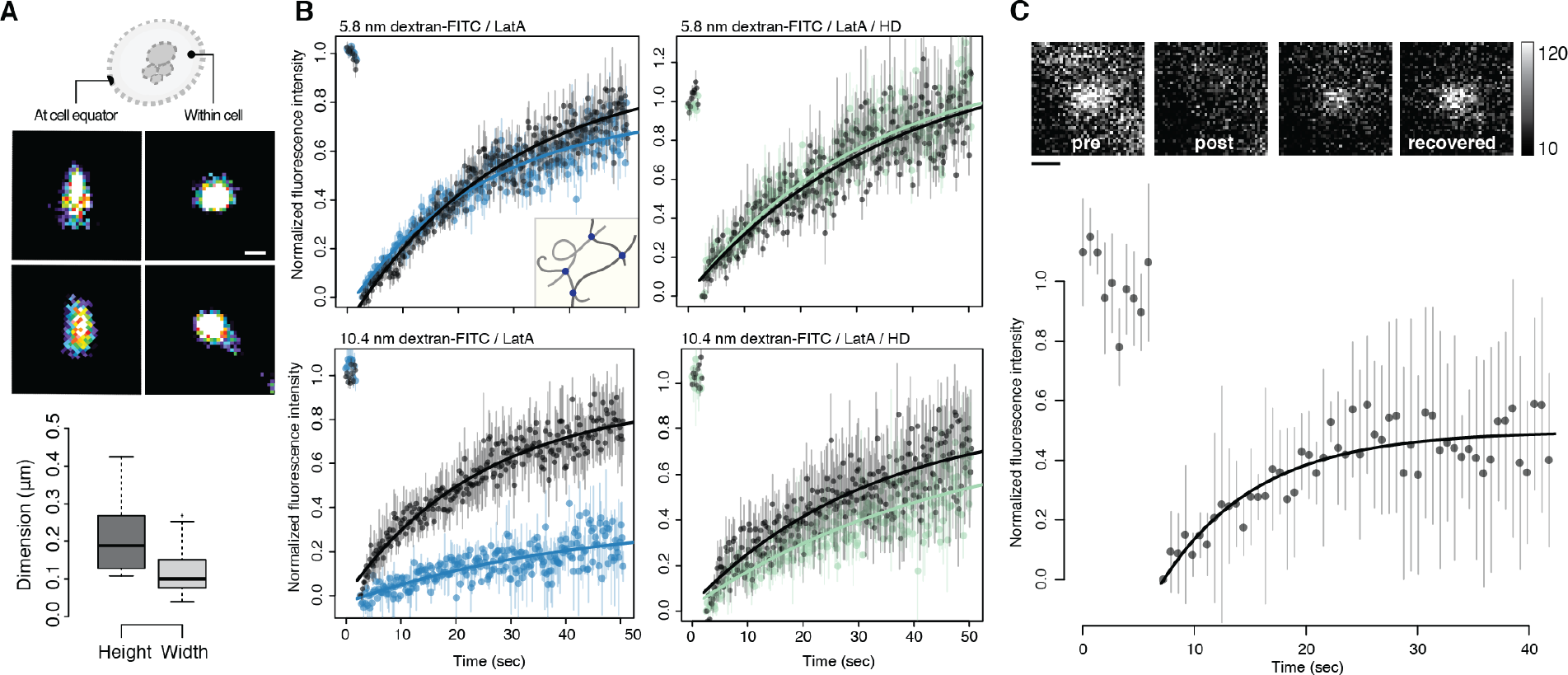
Endocytic puncta form dense condensates of labile molecules. (A) Dimensions of Sla1-GFP-labeled puncta measured using super-resolution microscopy (dSTORM). Lateral x, y resolution was ~10 nm. Pseudo-color reconstructed images show circular structures when viewed from the top, or within cells (top right), and narrow ellipses when imaged at the equator of cells (top left). Automatic segmentation (center) was performed on these images to determine the length (209 ± 10 nm) and width (118 ± 6 nm) of the endocytic puncta (n = 250). (B) Endocytic puncta are dense lattices that exclude molecules having dimensions > 10 nm. Fluorescence recovery after photobleaching (FRAP) of dextran-FITC molecules within endocytic puncta or neighbouring cytosol. FRAP of the bleached 5.8 nm dextran-FITC within either a Sla1-mCherry (top left panel; Lat A treated cells; blue) or a Syp1-mCherry (top right panel; Lat A and 5% HD treated cells; green) puncta and neighbouring cytosol regions (black) without Sla1 or Syp1 signals, respectively. Insert illustrates porous latticework composed of amorphous protein chains (grey filaments) with binding sites (dots) through which they are non-covalently associated. Same experiment with 10.4 nm dextran-FITC that scarcely permeate the Sla1 puncta (bottom left panel; Lat A treated cells; blue) but are mobile when puncta are dissolved by HD (bottom right panel; Lat A and 5% HD treated cells; green). Data points (mean ± SEM; n = 10 cells) were fitted to a single term recovery function (Methods). (C) Coat proteins exchange with endocytic puncta at rates typical of those observed for biomolecular condensate proteins. FRAP of Sla2-GFP: Signal recovery was measured within a segmented Sla1-mCherry region of interest to ensure that FRAP was acquired within the endocytic puncta (mean ± sd; n = 10 cells. Data fitted to a single term exponential, see Materials & Methods). Incomplete fluorescence recovery suggests that endocytic puncta are viscoelastic. Representative foci images (insert) before and upon bleaching and after recovery. 8-bit grayscale values, 10 to 120. Scale bar, 250 nm.

To determine whether endocytic condensates have a characteristic mesh size, we measured the diffusivity and permeability of probe molecules within and between condensates and the cytosol and at endocytic sites in cells where condensates were dissolved using 1, 6-hexanediol (HD). HD has been shown to dissolve a number of cellular structures, including different types of protein condensates *in vivo* (Rog, Köhler et al. 2017) and *in vitro* (Alberti, Saha et al. 2018). Sla1-labeled puncta as well as those labeled with all of the PLD-containing proteins that are associated with endocytic puncta, including Sla2, Ent1, Ent2, Yap1801 and Yap1802 dissolve in the presence of HD (*SI Appendix*, Fig. S3*A*) (Updike, Hachey et al. 2011, Kroschwald, Maharana et al. 2015). However, Syp1-mCherry, a membrane-bound endocytic initiator containing a scaffold-forming F-BAR domain, was insensitive to HD (*SI Appendix*, Fig. S3*B*). This suggests that membrane targeting of endocytic sites through F-BAR domain oligomerization was unperturbed by HD. We also noted that HD does not disrupt membrane integrity, as monitored by membrane leakage of carboxy-fluorescein (*SI Appendix*, Fig. S3 *C-E*). These results imply that HD disrupts endocytic condensates and therefore, in cells treated with HD, the diffusivity and permeability of endocytic sites should be identical to those of the surrounding cytosol.

We quantified the apparent mesh sizes within the endocytic puncta using Fluorescence Recovery After Photobleaching (FRAP) and colocalization of FITC-conjugated dextran molecules of 2.4, 5.8 and 10.4 nm dimensions. The dextran probes were osmoporated into cells and FRAP was measured within either Sla1- or Syp1-labeled puncta. Maintenance of cell mechanical properties following osmoporation was evaluated by monitoring passive diffusion of expressed viral microNS protein nanoparticles (*SI Appendix*, Fig. S7 *A-F*) (da Silva Pedrini, Dupont et al. 2014). Syp1 was used as a reference marker for cortical patches because it was not sensitive to HD at concentrations that dissolved endocytic puncta (*SI Appendix*, Fig. S3*B*). Although the density of the FITC-labelled 5.8 nm dextran was lower in Sla1-labeled puncta than in the surrounding cytosol, the mobility of the 5.8 nm probe was equivalent in both regions (Fig. 2B and *SI Appendix*, S4 *C-F*). Similarly, the fluorescence intensities of 2.4 nm and 5.8 nm FITC-labelled dextran recovered equally well in the Sla1-labeled puncta and cytosolic zones (Fig. 2*B* and *SI Appendix*, S4*F*). It is known that weakly charged molecules such as proteins do not encounter hindered diffusion across phase boundaries (Münchow, Schönfeld et al. 2008). Instead, the key determinants of diffusion of proteins across phase boundaries are the differential affinities to the coexisting phases and the concentration gradients (Münchow, Schönfeld et al. 2008), and our data are consistent with these expectations.

In contrast to the transport of 2.4 nm and 5.8 nm FITC-labelled dextran particles, only a few 10.4 nm dextran molecules were taken up by the endocytic puncta, although these were mobile in the cytosol (Fig. 2B). Furthermore, when Sla1-labeled puncta were disrupted by HD, we observed equivalent mobility of 10.4 nm FITC-labelled dextran molecules within cortical membrane sites, labelled with HD-resistant protein Syp1-mCherry, and the neighboring cytosol (Fig. 2B). Our results suggest that endocytic condensates are porous bodies with mesh sizes that are between 5-10 nm. These results are consistent with mesh sizes of networks of proteins that engage in non-covalent physical crosslinks in biomolecular condensates, and with electron microscopy data showing that endocytic puncta exclude ribosomes (Kukulski, Schorb et al. 2012, Wei, Elbaum-Garfinkle et al. 2017, Alberti, Saha et al. 2018).

To measure the exchange of endocytic proteins between condensates and the cytosol, we used FRAP to measure the dynamics of the coat protein Sla2. To control for dynamic Sla2 recruitment that occurs early in formation of the endocytic puncta, we evaluated fluorescence recovery during periods in which the apparent number of Sla2 molecules in the fluorescent foci of individual puncta was constant. FRAP measurements of entirely bleached whole puncta showed signal recovery within seconds and equivalent mobile and immobile fractions (0.50 ± 0.02; mean ± sem) for the protein Sla2, indicating that mobile fractions of Sla2 proteins are able to exchange with their surroundings on time scales of tens of seconds (Fig. 2*C*).

### PLDs are necessary for the formation of functional endocytic condensates

Next, we explored whether endocytic condensates formed by PLD-containing proteins require their PLDs to associate with the condensates and enable actin-independent endocytosis. Since HD generally disrupts interactions among PLD-containing proteins in biomolecular condensates we tested whether dissolution of Sla1-labeled puncta with HD inhibited both actin-dependent and independent endocytosis in *GPD1Δ* cells as quantified by the uptake of the external fluid-phase (Measured by Lucifer Yellow (LY) uptake) and membrane components (Measured by FM4-64 uptake), respectively (Fig. 3*A, B*). Dissolution by HD (concentration of 5% wt/v) led to inhibition of the uptake of membrane components. The HD concentration we used is considerably lower than the concentration necessary to disrupt membrane or Syp1-labeled cortical patch structures (Fig. 3*B*, *SI Appendix*, S3 *B-E*). Importantly, the inhibitory effect of HD on endocytosis was independent of turgor pressure or actin polymerization (Fig. 3*A* and *SI Appendix*, Fig. S5).

**Figure 3.**
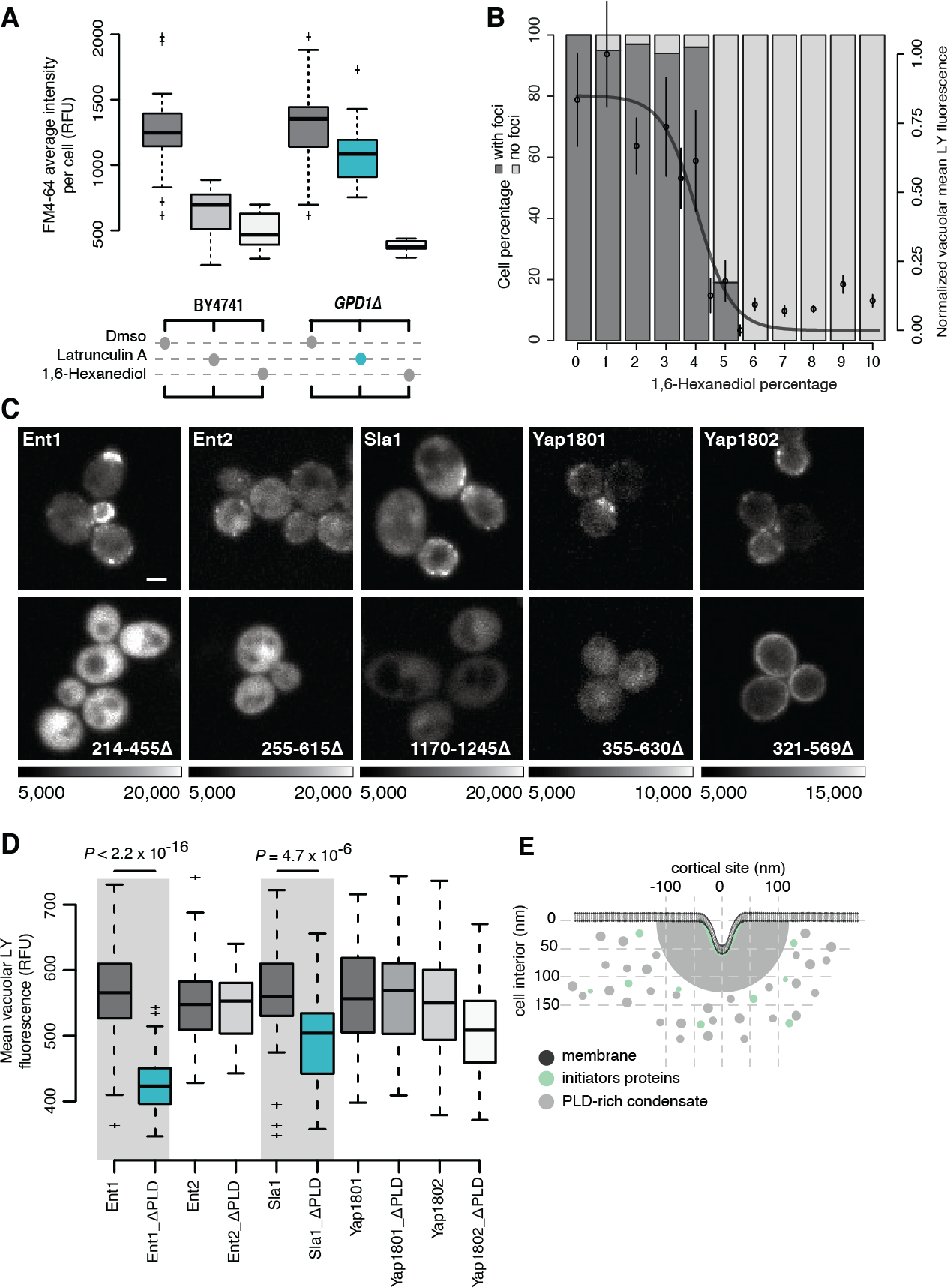
Puncta formed by PLD-containing proteins are necessary for actin-independent endocytosis. (A) Quantification of the membrane uptake of a lipophilic membrane-bound fluorescent dye (FM4-64) in *wildtype* BY4741 (left) and *GPD1*Δ cells (eliminates turgor pressure; right) treated with either DMSO, latrunculin A (prevents F-actin polymerization) or 1,6-hexanediol (Non-specifically alters solvent quality). Each boxplot shows the relative fluorescence units of n = 50 cells. *GPD1Δ* cells can undergo endocytosis in the absence of F-actin polymerization (blue) because there is no turgor pressure in these cells. (B) At fixed temperature, HD dissolves endocytic puncta and this in turn inhibits endocytosis. The bar plot quantifies the percentage of cells that contain Sla1-GFP foci (dark grey), or not (light grey), as a function of HD concentration monitored as counts of endocytic puncta labeled with Sla1-GFP (n = 150 cells). Plot overlay: fluid uptake of water-soluble fluorescent dye Lucifer Yellow (LY) into vacuoles (mean ± sd; n = 25 foci; logistic fit). (C) Prion-like domains (PLDs) of endocytic coat proteins are essential for their localization to endocytic puncta. Fluorescence images of cortical localization of Ent1, Ent2, Sla1, Yap1801 and Yap1802 fused to Venus YFP. Full-length (upper panels) *versus* C-terminal PLD truncation mutants of the proteins (lower panels). Sequence ranges of deleted PLDs indicated (lower) and grayscale dynamic ranges for image pairs. Scale bar, 2 μm. (D) Lucifer yellow dye uptake for strains that express either full-length or PLD deletion mutants of Ent1, Ent2, Yap1801, Yap1802 and Sla1 (as detailed in panel C). Reductions in Ent1 and Sla1 mutants are significant (n = 100 cells; two-sided t-test; see Materials and Methods). (E) Illustration of the membrane topology (dark grey) and remodelling into the cell during endocytosis in the absence of actin. Electron microscopy data suggest that clathrin-coated plasma membrane patches are surrounded by a cortical body of ~200 nm diameter (light grey) before appearance of actin structures (6). Clathrin heavy and light chains (Chc1 and Clc1) interact with initiator proteins (Ede1 and Syp1) to form a lattice on the membrane (in green). Subsequently, PLD-containing coat proteins (light grey), such as Sla1/2, Ent1/2, and Yap1801/2, directly bind to the initiator-clathrin lattice and form the endocytic puncta (in grey).

Next, we tested the specific importance of PLDs in endocytosis. For this, we tested whether specific PLDs are required for the assembly of endocytic condensates and actin-independent endocytosis. Deletion of the PLDs of Sla1 and Ent1 caused mis-localization of the Sla1, Ent1, Ent2, Yap1801 or Yap1802 proteins and also impaired endocytosis for Sla1 or Ent1 with their PLDs deleted (Fig. 3 *C-D*). These observations were consistent with previous reports regarding the effects of deletion mutants of Sla1 and Ent1 (Warren, Andrews et al. 2002, Mukherjee, Coon et al. 2009). Recruitment of Ent2, Yap1801 and Yap1802 to endocytic condensates required their PLDs but were not essential to endocytosis, though they may serve other roles in regulating endocytosis.

### The interaction network within endocytic puncta is governed by sequence features of PLDs

Recent studies have shown that multicomponent biomolecular condensates such as stress granules are governed by a core macromolecular network with an uneven distribution of interactions across nodes of the network (Sanders, Kedersha et al. 2020, Yang, Mathieu et al. 2020). Such networks are wired by specific types of network motifs or features. The functional importance of Sla1 and Ent1 and their PLDs suggests that they play the roles of scaffolds in organizing endocytic condensates (Kukulski, Schorb et al. 2012, Wei, Elbaum-Garfinkle et al. 2017, Alberti, Saha et al. 2018). To probe the organization of the protein-protein interaction network that underlies endocytic condensates, we performed an *in vivo* screen for protein-protein interactions. Specifically, we used a Protein-fragment Complementation Assay with the reporter protein DiHydroFolate Reductase (DHFR PCA) to quantify the effects of PLD deletion on *in vivo* interactions involving or mediated by Sla1 (Fig. 4*A*). Selection for DHFR reconstitution with Sla1 in which the PLD was deleted (*Sla1* Δ*PLD*) revealed the preservation of only two out of the thirteen interactions that were detected with the full length Sla1. The interactions that were preserved are relevant only to actin-*dependent* endocytosis (Fig. 4 *B-D*). These results reveal sequence features of the Sla1 PLD that coordinate the network of interactions and organization of endocytic condensates.

**Figure 4.**
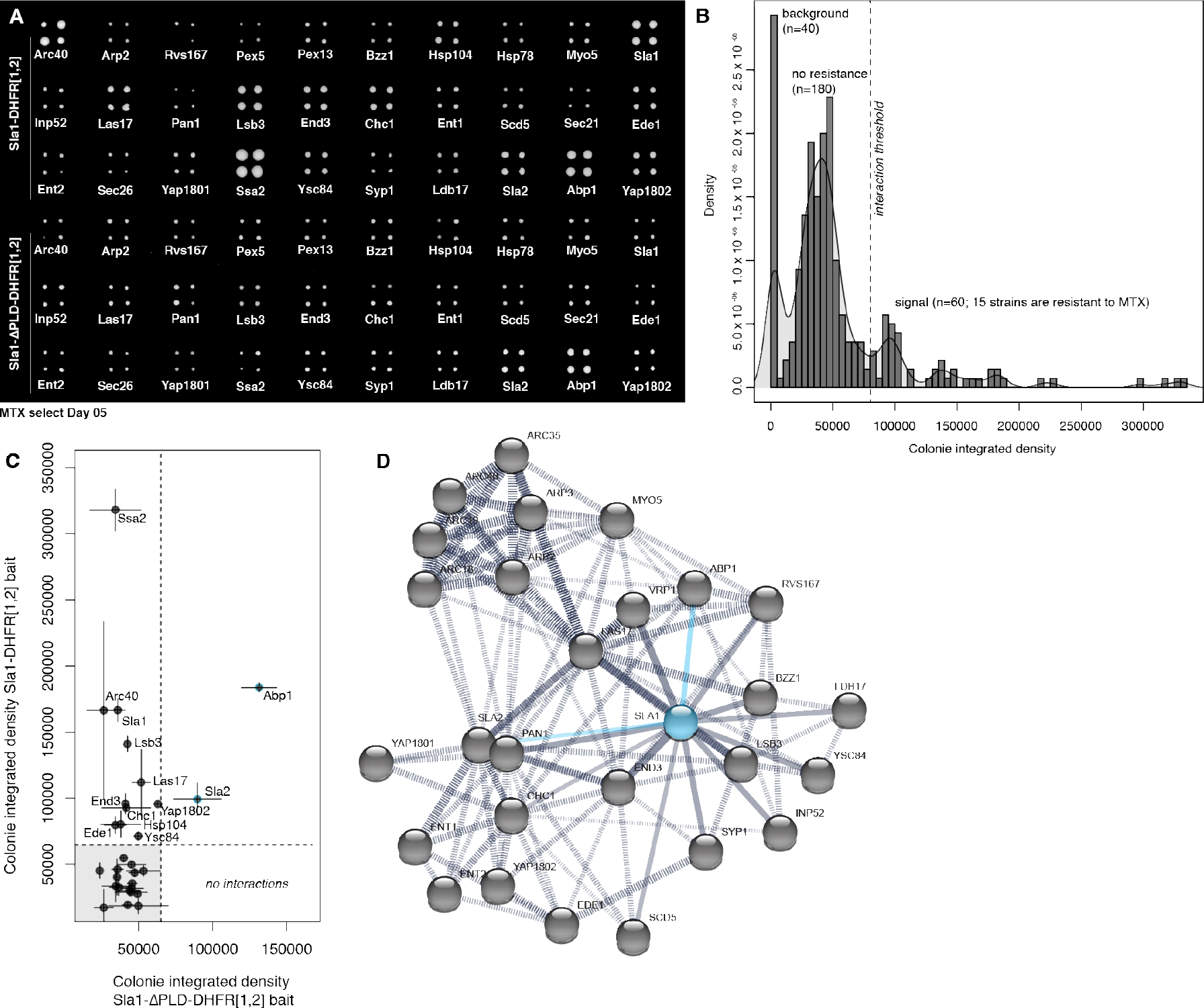
Proteins can partition into endocytic puncta through a network for interactions coordinated by PLDs. (A) DHFR-PCA and methotrexate (MTX) selection plate after 5 days of growth shows that the PLD deletion mutant of Sla1 loses all but two interactions of *wildtype* Sla1. (B) Density (number of colonies with given total pixel intensity) *versus* colony integrated growth (total pixel intensity) after 5 days of MTX selection for DHFR reconstitution. Peaks are labelled with their corresponding measurements for plate background (n=40 empty spots), colonies with no resistance to MTX (n=180 colonies; 45 strains) and colonies that grow on MTX (n=60 colonies; 15 strains) above a selected integrated density threshold. We detected 13 interactions for full-length Sla1 protein. (C) Colony integrated growth (total pixel intensity) after 5 days of MTX selection for DHFR reconstitution with *wildtype* Sla1 *versus* colony integrated growth (total pixel intensity) after 5 days of MTX selection for DHFR reconstitution with PLD deletion mutant of Sla1. Compared to *wildtype* Sla1 interactions, we detect 2 interactions (blue points) for the mutant Sla1 protein that do not contain the PLD. (D) Protein interaction network from STRING (version 11.0) for the selected subarray of 30 potential interactors (grey dots) of Sla1 (blue dot). We represent the 13 direct Sla1 interactions (solid black lines) and DHFR-PCA interactions with Apb1 and Sla2 that are preserved with the PLD deletion mutant of Sla1 (solid blue lines) amongst all the protein-protein interactions (dashed black lines) determined experimentally.

### Glutamine-rich PLDs are required for proteins to partition into endocytic puncta

Given that the PLD of Sla1 is central to coordinating the network of essential protein-protein interactions, we reasoned that determinants of protein partitioning into endocytic condensates and the interactions among them should depend on the amino acid compositions of their PLDs (Patel, Lee et al. 2015, Boke, Ruer et al. 2016, Alberti, Saha et al. 2018, Wang, Choi et al. 2018, Vijayakumar, Perrois et al. 2019). PLDs have distinctive compositional biases that include ratios of Asn *versus* Gln and the uniform sequence distribution of aromatic amino acids (Martin, Holehouse et al. 2020). Alberti *et al*, showed that PLDs that drive the formation of amorphous and insoluble amyloid fibrous bodies have a clear preference for Asn over Gln. Conversely, PLDs with higher Gln:Asn ratios form soluble puncta (Alberti, Halfmann et al. 2009). Sla1, Sla2, and the endocytic epsin proteins Ent1 and Ent2 have high Gln and Asn contents with a clear preference for Gln. The Gln:Asn ratios in these PLDs are 2:1 in Sla1, ~5:1 in Sla2, ~3:1 in Ent1, and ~4:1 in Ent2, respectively.

To test if higher Gln content is essential for partitioning into and the material properties of endocytic condensates, we used artificial PLDs to substitute for the native endocytic PLDs. Specifically, we leveraged the work of Halfmann *et al*., who compared the prion-forming potential of the NM domain of *wildtype* Sup35 (Sup35 NM), which includes a PLD, to those of Sup35 in which all NM domain Gln residues are substituted for Asn (Sup35 NM (N)) or all Asn substituted with Gln (Sup35 NM (Q)) (Halfmann, Alberti et al. 2011). The Gln:Asn ratios in these PLDs are ~1.5:1 (Sup35 NM), 1:33 (Sup35 NM (N)), and 9:1 (Sup35 NM (Q)). We designed different chimeras of the endocytic epsin proteins Ent1 and Ent2, in which the cognate PLDs within these proteins were replaced with either that of Sla1 or variants of the non-endocytic protein Sup35 NM domain (Fig. 5*A*). Although, the Sla1 PLD shares little sequence identity with either Ent1 or Ent2 PLDs (*SI Appendix*, Fig. S6*A, B*), Sla1 PLD chimeras of Ent1 and Ent2 partitioned into endocytic condensates (Fig. 5*B, C* and S6 *C-G*). Lowering the Gln:Asn ratio or inverting this in favor of Asn increases the amyloid forming potential of PLDs (Halfmann, Alberti et al. 2011). We tested additional Ent1 or Ent2 chimeras wherein the cognate PLDs were replaced with Sup35 NM, NM(N) or NM(Q). We assessed the effects of these substitutions on the partitioning of chimeras into endocytic condensates (Fig. 5*A*). Consistent with the need for a clear asymmetry between the numbers of Gln *versus* Asn, we observed that only the Sup35 NM(Q) chimeras of Ent1 and Ent2 partitioned into Sla1-labeled endocytic condensates (Fig. 5*E*, *SI Appendix*, S6*F, G*). Conversely, the Sup35 NM and Sup35 NM(N) chimeras did not partition into endocytic condensates; instead, they formed distinct puncta, indicative of structural features and interaction preferences that are distinct from those of endocytic condensates (Fig. 5*E, G*). We further surmised that the fibrillar content of Sla1 labeled puncta must be low; to test this hypothesis, we measured the staining of Sla1 labeled endocytic condensates using Thioflavin (ThT), which is a marker for amyloid-like aggregates (*SI Appendix*, Fig. S6*H, I*) (Alberti, Saha et al. 2018, Wang, Choi et al. 2018). These data showed an absence of ThT staining in Sla1-labeled endocytic condensates or in puncta formed by Sup35 NM(Q) (Halfmann, Alberti et al. 2011).

**Figure 5.**
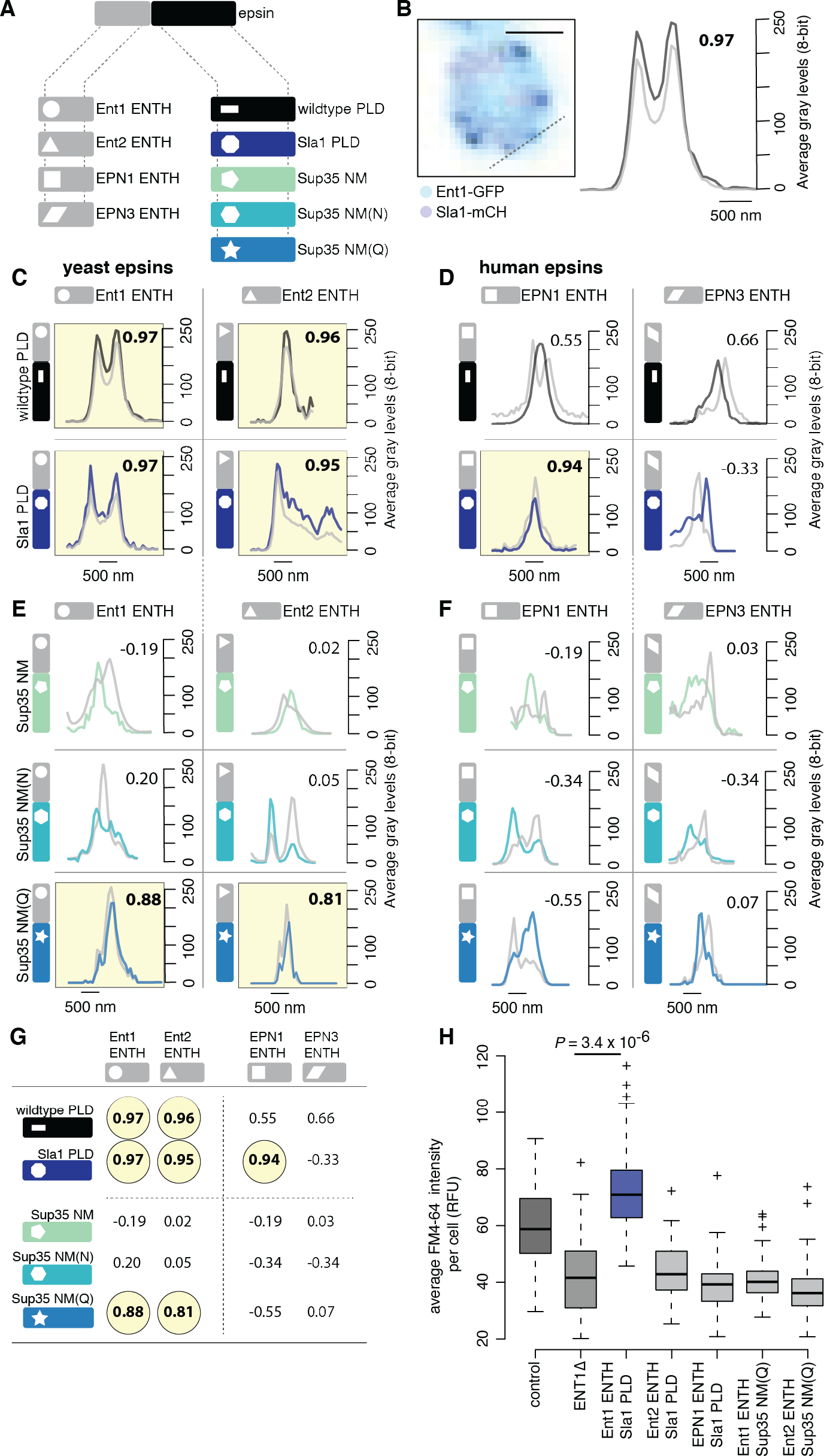
PLDs with shared sequence features can be interoperable and drive assembly of functional endocytic condensates. (A) Illustration of the general structure of epsin proteins (top) with the four ENTH modules (bottom left; shape coded) and five PLD modules (bottom right; colour and shape coded) we used to construct sixteen different protein chimeras. (B) Colocalization of Ent1-GFP signal (black) with signal of Sla1-mCherry labelled endocytic puncta (gray) in yeast cells (left), line scan on confocal fluorescence images (right) was performed as indicated on the cell image (left; dashed line). Pearson correlation values between the fluorescence signals are given (top right of plot) to confirm whether the signals colocalize (values above 0.8 are considered significant and highlighted in yellow in panels C-F; corresponds to a *p-value* < 0.005). Scale bar 1 μM. (C) Wildtype yeast epsin proteins Ent1 and Ent2 (top) and chimeras of Ent1 ENTH and Ent2 ENTH respectively fused to Sla1 PLD (bottom). (D) Wildtype human epsin proteins EPN1 and EPN3 (top) and chimeras of EPN1 ENTH and EPN3 ENTH respectively fused to Sla1 PLD (bottom). (E) Chimeras of Ent1 ENTH and Ent2 ENTH respectively fused to either Sup35 NM (top), Sup35 NM(N) (middle) or Sup35 NM(Q) (bottom). (F) Chimeras of human EPN1 ENTH and EPN3 ENTH respectively fused to either Sup35 NM (top), Sup35 NM(N) (middle) or Sup35 NM(Q) (bottom). (G) Summary of the Pearson correlation values for colocalization of all wildtype and chimera proteins with endocytic puncta described in C-F with. Values above 0.8 are considered significant and highlighted in yellow, corresponding to a *p-value* < 0.005. (H) FM4-64 dye uptake in cells for strains that overexpress fusions of the yeast Ent1 ENTH, yeast Ent2 ENTH or human EPN1 ENTH respectively fused to either Sla1 PLD or Sup35 NM(Q) that are shown to colocalize with endocytic puncta in C-F (n = 100 cells; two-sided t-test; see *SI Appendix, Extended Materials and Methods*).

Our data show that the compositional features of PLDs, particularly the asymmetry between numbers of Gln *versus* Asn, are essential for defining the selective partitioning of proteins into endocytic condensates. In addition, the human orthologs of epsins, EPN1 and EPN3 expressed in yeast did not partition into Sla1-labeled endocytic puncta (Fig. 5*D* and *SI Appendix*, S6 *C-E*). Among the human chimeric proteins that we generated, only the Sla1 PLD chimera of EPN1 partitioned to endocytic puncta, whereas all other chimeras did form independent puncta (Fig. 5*D, F, G*). These data point to a clear preference for Gln-rich, non-amyloid forming PLDs derived from yeast, which have different compositional biases than proteins with PLDs from other organisms. It is also worth noting that even if several PLDs share generic physical properties that result in their partitioning into the same biomolecular condensates, each resulting condensate does not have the same material properties (Wang, Choi et al. 2018). This is important because material properties of endocytic condensates could tune to their mechano-active potentials. Consistent with this view, among all of the PLD chimeras of Ent1, including those that did partition to endocytic condensates, only the Sla1 chimera supported endocytosis as measured by FM4-64 dye uptake in *ENT1Δ* cells (Fig. 5*H*).

## Discussion

We have presented data to show that endocytic condensates form at cortical sites and these hemispheric bodies require cohesive interactions provided by PLDs within endocytic coat proteins. Endocytic condensates form reversibly, and they are required to drive clathrin- and actin-independent endocytosis. How might condensates contribute to endocytosis in the absence of actin? Several mechanisms could act in synergy with the formation of the endocytic condensates to drive mechanical invagination of the plasma membrane. These include (a), contributions from membrane curvature-inducing proteins and protein complexes, including convex-shaped BAR (for Bin, Amphiphysin and Rvs) domain-containing proteins (Youn, Friesen et al. 2010, Yu and Schulten), (b) insertion of an amphipathic helix into the outer leaflet of the membrane bilayer, which pushes the head groups apart (Ford, Mills et al. 2002, Boucrot, Pick et al. 2012), (c) modulation of lipid composition (Graham and Kozlov 2010, Anitei, Stange et al. 2017), (d) local relief of turgor pressure (Scher-Zagier and Carlsson 2016) and (e) “steric pressure” exerted at cortical sites due the excluded volumes of proteins that encompass certain categories of disordered domains (Busch, Houser et al. 2015, Snead, Hayden et al. 2017). Finally, analogous to condensates formed by synthetic polymers that are known to drive membrane vesicle formation from artificial phospholipid bilayers (Li, Lipowsky et al. 2011), we postulate that cohesive interactions that contribute to the formation of endocytic condensates also make these condensates mechanoactive by providing the free energy to drive membrane remodeling (Bergeron-Sandoval and Michnick 2018).

Of the mechanisms enumerated above, two stand out for the distinctive roles that are ascribed to intrinsically disordered regions (IDRs) of proteins, the steric pressure model and our mechanoactive model (Bergeron-Sandoval and Michnick 2018, Yuan, Alimohamadi et al. 2020). Stachowiak and coworkers have proposed that IDRs can drive membrane remodeling and endocytosis primarily via excluded volume effects – a phenomenon they refer to as “steric pressure” (Busch, Houser et al. 2015). Their recent studies also suggest that phase separation of IDRs involved in generating steric pressure also undergo phase separation via depletion-mediated forces (Yuan, Alimohamadi et al. 2020). Of course, not all IDRs adopt expanded conformations with high excluded volumes (Das, Ruff et al. 2015). Instead, IDRs come in different flavors; for instance IDRs such as the PLDs of Sla1, Sla2, Ent1, and Ent2 are deficient in charged or proline residues and are characterized by cohesive interactions with one another leading to compact conformations with average radii similar to those of folded proteins (Ruff, Pappu et al. 2019). Therefore, there appear to be two distinct categories of IDRs that drive membrane invagination namely those that do so *via* “hard” repulsive interactions afforded by expanded IDRs and a different category of IDRs that drive condensate formation *via* cohesive interactions, as occurs in our mechanoactive model (Bergeron-Sandoval and Michnick 2018). This raises the interesting prospect of synergies between the two curvature generation mechanisms by two distinct classes of IDRs.

We hypothesize that the energy stored with endocytic condensates can be converted into mechanical work to deform the membrane and the cytosol. The mechanics of this process can be described by analogy to a soft viscoelastic and sticky balloon bound to a soft elastic sheet (Movie S1). A balance between binding and the elastic/surface deformation energies is achieved upon membrane invagination. This idea can be captured in a simple phenomenological model where we express the mean-field energy *U* stored in a condensate as the sum of mechanical strain energy (ϕ term) and work (ψ term), respectively (*SI Appendix*):

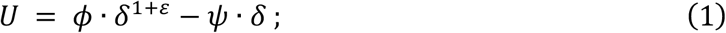

Here, *U* is a mean-field energy, δ is the invagination depth of both the membrane and cytosol (which are coupled by virtue of conservation of volume of the condensate) and the exponent ε > 0 is determined by the deformation geometry (*SI Appendix*). At equilibrium, (∂*U*/∂δ) = 0, and we expect invagination to balance the two contributions such that the value of δ* that minimizes the mean-field energy (1) is computed as,

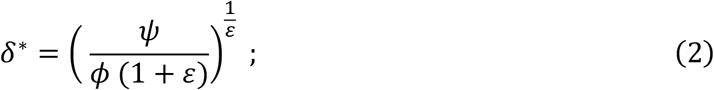

Equation (2) shows that the invagination depth δ is determined by the ratio ψ/ϕ and the deformation geometry that is captured in the exponent ε. The numerical values of ϕ and ψ can be estimated from the dimensions of the condensates as well as the viscoelastic properties of the cytosol, the condensate, and the membrane, respectively (*SI Appendix*).

To test whether the endocytic condensates have viscoelastic properties that are required to drive membrane invagination we used active rheology to measure the material properties of the cytosol in which endocytic condensates are embedded. Since the endocytic condensates are too small to probe their material properties directly, we inferred their material properties using the contact theory of Hertz (Hertz 1882). This theory relates the elastic moduli of elastic materials in contact with one another through the resulting geometries of their contacting surfaces. We probed the material properties of the yeast cytoplasm with optical tweezers to measure the frequency-dependent amplitude and phase responses of 200 nm diameter polystyrene beads that are embedded in cells (Fig. 6, *SI Appendix*, Fig. S7*G, H*). We used an acousto-optic device to oscillate the position of the optical trap in the specimen plane at frequencies that spanned over four orders of magnitude and measured the displacement of trapped beads from the trap center using back focal plane interferometry (Fig. 6 *A-C*). We quantified the viscoelastic properties of the cytosol surrounding the beads by measuring the phase and amplitude of displacements of beads in response to oscillations of the optical tweezers and calculated the power spectrum of unforced fluctuations of the bead to obtain storage (G’) and loss (G”) moduli as a function of frequency (Fig. 6*B, C* and *SI Appendix*) (Hendricks and Goldman 2017). We used both the dimensions and time-lapse fluorescence imaging of Sla1-labeled puncta to determine that endocytic condensates expand at a rate of 2360 ± 120 nm s^-1^ (*SI Appendix*, Fig. S7*I, J*), corresponding to a stress at ~30 ± 2 Hz. At this timescale, the cytosol is principally elastic with a shear modulus of approximately 20 Pa (Fig. 6*C*). We also measured the linear displacement of Sla1-labeled puncta within the confocal volume as a function of time. Membrane invagination occurs at a velocity of 7.4 ± 2.5 nm s^-1^, and this corresponds to a frequency of 0.1 ± 0.04 Hz (*SI Appendix*, Fig. S7*I, J*). In this regime, the cytoplasm is viscoelastic and mechanically compliant (Fig. 6*C*). The G’ and G” values we measured here are similar to those of the cytoplasm of adherent mammalian cells (Hendricks, Holzbaur et al. 2012, Guo, Ehrlicher et al. 2013, Guo, Ehrlicher et al. 2014).

**Figure 6.**
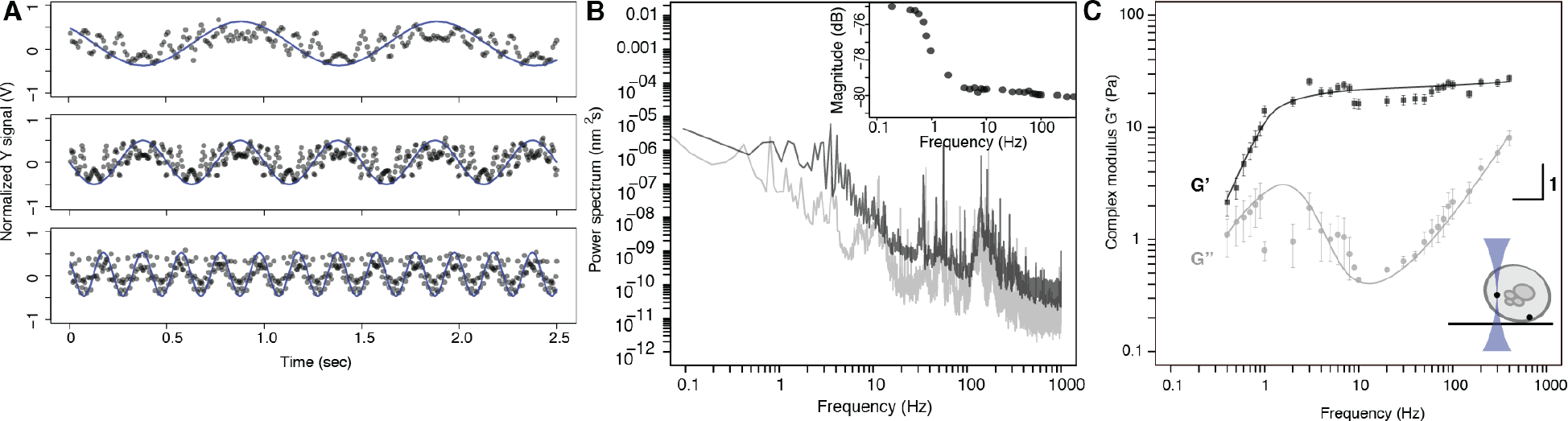
Sla1-labelled condensates are embedded in a viscoelastic cytosol. (D) Position sensitive detector (PSD) output signal in volts (V) *versus* time for acquisitions made at 1Hz (top), 2 Hz (middle) and 5 Hz (bottom). A bead located in the cytoplasm was oscillated with the AOD in the Y-axis of the specimen plane with fixed tweezer movement amplitude (normalized blue curve) at different frequencies. The recorded PSD raw traces (black points) were normalized to a corresponding magnitude range (coherence cutoff of 0.9). (B) Power spectrum of the oscillated bead (black) with magnitude of response as a function of frequency (insert). (C) Decomposition of G* as a function of frequency into G’ (storage modulus; darker squares) and G” (loss modulus; light shade circles) for beads distributed at both the cell periphery and interior (see schematic insert; mean ± sd; n = 17 cells) with an average trap stiffness k_trap_ (mean ± se; 8.0 x 10^-5^ ± 2.7 x 10^-5^ N m^-1^) and photodiode sensitivity factor *β* (mean ± se; 10.7 x 10^3^ ± 2.3 x 10^3^ nm V^-1^).

Based on the parameters obtained from active rheology including the dynamic modulus of the cytoplasm, we were able to determine the material properties of endocytic condensates embedded in the cytoplasm using Hertz theory and measurements of the dimensions of condensates that we determined by super-resolution microscopy (Fig. 2*A*) (Hertz 1882). We used this information to estimate the apparent Young’s modulus of endocytic condensates to be 59 Pa, which is on the same order as that of the cytosol at 45 Pa at 1 Hz (Fig. 6*C*, and *SI Appendix, Extended Materials and Methods; Eq. 3.7-3.10*). These results are consistent with those obtained for other protein-based elastic materials (Reichheld, Muiznieks et al. 2017).

The Young-Laplace equation yields an estimate of γ_dc_ = 7 × 10^-5^ N•m^-1^ for the interfacial tension at the condensate-cytosol interface. This estimate is based on the pressure difference across the cytosolic interface and the mean curvature of the condensate (*SI Appendix, Extended Materials and Methods; Eq. 4.6*). It falls within the range of values that have been reported for other protein condensates, including nucleoli and P granules (*SI Appendix*, *Extended Materials and Methods; Eq. 4.9*) (Brangwynne, Mitchison et al. 2011, Elbaum-Garfinkle, Kim et al. 2015).

Using the inferred apparent elastic modulus for the endocytic condensate, we computed the mechanical strain (φ) and mechanical work (ψ), respectively as functions of membrane and cytosol invagination δ (Fig. 7*A*, *SI Appendix*, S8 and *Extended Materials and Methods; Eq. 4.25, 4.26*). Our physical framework suggests that the formation of endocytic condensates generates 4.9 × 10^-18^ J (Fig. 7*A* and Table S*4*), which is within the range of the energy required to provide the necessary membrane invagination. Rheology experiments also yield estimates for the energy of deformation that are strikingly similar. This points to a clear route for force generation by condensates that enables membrane invagination, thereby activating membrane constriction and vesicle scission to drive actin-independent endocytosis (Idrissi, Blasco et al. 2012).

**Figure 7.**
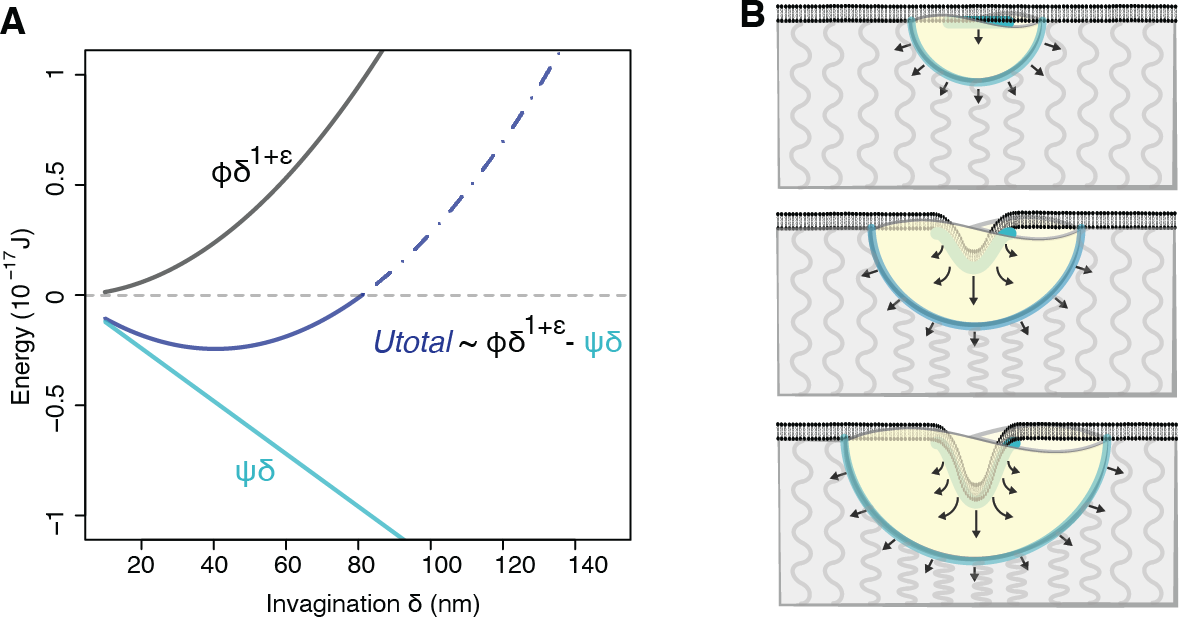
Endocytic condensates do mechanical work to deform the membrane and cytosol. (A) Energy U_total_ *versus* membrane invagination d determined from equation (1), (inset) determined by opposing (ϕ) and favoring (ψ) energy terms. *U*_total_ (dark blue), ϕ (black) and ψ (light blue) versus δ. The energy is favourable for δ between about 15-80 nm (solid blue line) and unfavorable above 80 nm (dashed blue line). Quantities used to calculate energies are detailed in Fig. S8*A*, Tables S2, S3 and S4. (B) Mechanical description of endocytic condensate-driven membrane invagination. Endocytic condensate (yellow) binds to (wets) the bilayer membrane (black) and drives membrane invagination as the condensate expands to maximize contact with the cytosol (top to bottom). Forces balance under a Young-Dupré adhesion gradient (blue lines and arrows) and resistance of the cytosol (grey curved lines).

A mechanical analogy can aid in understanding our model (Fig. 7*B*, *SI Appendix*, Movie S1). Adhesion of the endocytic condensate to the cytosol tends to drive the condensate to maximize the contact surface area between the condensate and the cytosol. Because the condensate is a viscoelastic material, its volume must remain constant. The area of the condensate-cytosol interface is proportional to the square of the radius. Any increase in interfacial area must be compensated by a decrease in volume of the condensate, which varies by the cube of the radius of the condensate. Since the drive to increase the condensate-cytosol contact is symmetrical, the only way that the condensate volume can be reduced, is by invagination of the membrane. This, however, increases the favorable condensate-membrane surface contact, thus acting as a positive feedback to further drive membrane invagination. The system comes to equilibrium when the viscoelastic properties of the condensate and cytosol prevent further expansion and invagination. This is the point at which scission of a mature endocytic vesicle must occur.

There remains the question of how the geometry of the observed membrane invagination comes about. Evidence from electron and super-resolution fluorescence microscopy indicates that the favored geometry of the membrane is flat with invagination centered in the middle of the endocytic condensate. Such geometries can be explained by a local radial stress gradient generated by adhesion of the viscoelastic condensate to the membrane on one side and the cytosol or the other (Korchagin, Dolbow et al. 2007). Local radial stress gradients can also be generated by asymmetries in local binding of adaptor proteins, or by distinct lipid compositions. Overall, our model and the questions they raise provide the motivation for continued investigation of the details of regulation of endocytosis in eukaryotic cells. Our model also provides motivation for investigating the mechano-active roles of viscoelastic condensates that contribute to vesicle trafficking processes and involve proteins with PLDs (Pietrosemoli, Pancsa et al. 2013).

## Acknowledgements

The authors acknowledge support from CIHR grants MOP-GMX-152556 (SWM), the US National Institutes of Health grant R01NS056114 (RVP), the Fonds Québécois de la Recherche sur la Nature et les Technologies (SWM and PF), the US National Science Foundation through grant MCB-1614766 (RVP), NSERC RGPIN/05843-2014 (AJE), CIHR 143327 (AJE), CFI 32749(AJE and AGH), and the Human Frontier Science Program RGP0034/2017 (SWM and RVP). CEC was supported by the National Institute of General Medical Sciences of the National Institutes of Health (NIH) under award T32GM008268. Research in the Keller Lab is supported by National Science Foundation award MCB-1402059 to SLK. We thank Cliff Brangwynne, Julien Berro, David Drubin, Alex Holehouse, Tom Pollard, and Kiersten Ruff for thoughtful discussions and advice on the manuscript; Jackie Vogel for strains; Susan Liebman, Simon Alberti and Randal Halfmann for plasmids; Jacqueline Kowarzyk and Philippe Garneau for technical assistance and Rosa Kaviani for help with FRAP experiments.

## Author Contributions

LPBS and SWM designed the research with the assistance of RVP; LPBS performed biological research; LPBS, AJE, RVP and SWM analysed the biological data; CC, CEC and SLK performed and analysed the vesicle leakage experiments; LPBS and HKH performed micro rheology experiments; LPBS, HKH, AJE and AGH analysed micro rheology data; LPBS, HKH and PF developed physical model; LPBS and SWM wrote the first version, all authors corrected the paper.

## Supplementary Materials

Materials and Methods

Figures S1-S6

Tables S1-S4

Movies S1-S2

