## Supplementary Materials for "Proteins with prion-like domains can form viscoelastic condensates that enable membrane remodeling and endocytosis"

**This PDF file includes:**

Materials and Methods

Figures S1 to S8

Tables S1 to S4

Captions for Movies S1

References

**Other Supplementary Materials for this manuscript includes the following:**

Movies S1

#### Materials and Methods

##### Strains and model system

We performed *in vivo* experiments in the *Saccharomyces cerevisiae* BY4741 *MATa his3Δ1 leu2Δ0 met15Δ0 ura3Δ0* background. To decouple the effects of turgor pressure on clathrin-mediated endocytosis (CME) we performed many experiments in a yeast strain with the gene for glycerol-3-phosphate dehydrogenase deleted, *GPD1Δ*. We acquired the *GPD1Δ* and other deletion strains from the Yeast Knock-out (YKO) deletion collection and GFP fluorescent strains from the Yeast GFP Clone Collection (Huh, Falvo et al. 2003), were generous gifts from J Vogel at McGill University. A complete list of strains used in this study are in Table S1.

##### Cell culture, gene manipulations and fluorescent reporters

Cells were normally grown to exponential phase ($OD_{600}$ 0.1 to 0.6) in either rich medium (YPD) or low fluorescence medium (LFM) (Sheff and Thorn 2004). Liquid media or solid-agar cultures were incubated at 30°C. For 2-color imaging and other specific needs, the coding sequences for different fluorescent proteins were integrated into the genome, 3' to reporter protein open reading frames (ORFs) by homologous recombination (Tarassov, Messier et al. 2008). In short, mCherry and Venus YFP tags were integrated *via* homologous recombination by amplifying the HPH or NAT resistance cassettes from the respective pAG32 or pAG25 vectors with primer tails homologous to flanking sequences to the respective loci. BY4741, *GPD1Δ* or other YKO strains were then transformed with the respective PCR cassettes, selected for HPH or NAT resistance in YPD medium and confirmed by diagnostic PCR.

##### Truncation and site-directed mutagenesis within ORFs

Integration of the coding sequence for fluorescent protein Venus YFP by homologous recombination into the genome was also used to truncate the prion-like domains (PLDs) sequences in the ORFs of Sla1, Ent1, Ent2, Yap1801 and Yap1802. In order to truncate the required gene fragments, the Venus YFP coding sequence was integrated at precise locations prior to the STOP codon. The amino acid sequences that are deleted in each PLD truncation mutant are; Sla1 (1170-1245*Δ*), Ent1 (214-455*Δ*), Ent2 (255-615*Δ*), Yap1801 (355-630*Δ*) and Yap1802 (321-569*Δ*).

To generate proline substitution point mutations within or near PLDs of Sla1 and Ent2; we cloned synthetic gene fragments (GenScript) that coded for the mutations into the respective MoBy-ORF p5472 plasmids of our target genes (Ho, Magtanong et al. 2009). The respective position of proline mutations, gene fragment size and restriction enzymes are; Sla1 G1224P, Q1227P, A1231P, F1234P (842 bp, NarI and XmaI); Ent2 R359P, Q362P, H366P, L369P, L385P, L388P, K401P, E404P, L408P, Q411P, L415P, Q418P (1395 bp, XhoI and AflII). Standard cloning procedures with matching restriction enzymes were used to build the constructs and we confirmed final constructs by sequencing. Respective deletion strains were then complemented with either the proline-mutated or original p5472 plasmids.

**Diffraction-limited fluorescence microscopy**

For most experiments, cells were grown in LFM to an $OD_{600}$ of ~0.1 to 0.6 and plated on either Nunc glass bottom 96-well plates (Thermo Scientific; 164588), glass bottom 8-well plates (Ibidi) or glass bottom 35 mm round dishes (MatTek). We used concanavalin A (Sigma-Aldrich ConA # C-7275) as a cell surface binding agent. Each well was loaded with 1mg/ml final concentration of ConA solution at room temperature for 15 minutes. ConA was then removed and wells were completely air dried before cells were added. Fluorescence images were acquired with distinct imaging platforms;

For the assessment of PLD truncations, we imaged cells on a Nikon TE2000 inverted microscope equipped with a 100X/1.45 plan APO lambda oil objective (Nikon), X-Cite lamp source (Excelitas), respective FITC (Chroma 41001HQ), EYFP (Chroma 49003ET) and mCH/TR (Chroma 49008ET) dichroic cubes and a Cool SNAP HQ camera. Z-stacks were acquired through a micron deep region with 5 planes and presented by maximal Z-projection.

For the measurements of lucifer yellow fluorescent probe uptake and cell sizes under different osmotic pressures, fluorescence images were collected on an InCell 6000 automated confocal microscope configured with a 100x/0.9 Plan FLUOR objective (Nikon) and 488 nm laser diode and FITC 525/20 emission filter for GFP fluorescence or 561 nm DPSS laser and dsRed 605/52 emission filter (GE healthcare life sciences). Single or two-color images were collected sequentially on a single focal plane with an exposure time of 100 msec. and a confocal slit of 2 AU. Image analysis and signal automated segmentation was performed with the InCell Developer software (GE healthcare life sciences) and the data was further analyzed and plotted in the R environment.

For other imaging data, the Quorum Diskovery platform was used in widefield, confocal and super-resolution imaging modes. Our Quorum Diskovery platform consists of a Leica DMi6000 inverted microscope equipped with a Diskovery multi-modal imaging system (Spectral) attached to either a Hamamatsu EM X2 camera or ORCA FLASH 4.0 V2 digital CMOS camera. Wide field or confocal excitation are achieved with a Spectral laser merge module with mounted 405 nm, 440 nm, 488 nm, 561 nm, and 640 nm diode pumped solid state laser sources linked to a Borealis beam conditioning unit. Images were acquired with a HCX PL APO 63x /1.47 NA oil corrected TIRF objective (Leica). This platform was remote controlled by the Metamorph software (Molecular Devices) and images were acquired and analyzed trough distinct pipelines. For particle tracking and mean squared displacement analyses, we used the Wave Tracer plugin to localize fluorescent foci centroid position through a wavelet algorithm and tracks particles in times stacks to calculate particles movement.

**Fluorescent probes to quantify endocytosis or detect amyloid structures**

Lucifer yellow (LY; Life technologies) assays to quantify endocytosis were performed at a final concentration of 1 mg/ml in YPD medium. Cells were incubated with the LY for 20 min or more. We then centrifuged at 3000 x g and washed cells 3 times in phosphate buffered saline (PBS; 137 mM sodium chloride, 10 mM phosphate, 2.7mM potassium chloride) before imaging in PBS with excitation wavelength ($\lambda_{ex}$) of 428 nm and emission wavelength ($\lambda_{em}$) of 536 nm. Measurements were taken from multiple cells in a single sample and this experiment was replicated 6 times.

The lipophlic stryryl dye FM4-64 (*N*-(3-Triethylammoniumpropyl)-4-(6-(4-(Diethylamino) Phenyl) Hexatrienyl) Pyridinium Dibromide) (Life technologies) was used to monitor plasma membrane uptake and staining of vacuolar membranes. Plasma membrane was labeled with 10 to 20 $\mu$M FM4-64 in YPD media and cells were incubated for 5 to 120 min. Cells were washed once in PBS and resuspended in LFM for imaging. FM4-64 stained cells were quantified by fluorescence microscopy ($\lambda_{ex}$ of 510 nm, $\lambda_{em}$ of 750 nm) on our Quorum platform (see Diffraction-limited fluorescence microscopy section).

We determined whether Sla1-mCherry puncta were labeled with the amyloid binding dye thioflavin T (ThT) in both live and fixed cells. Live cell ThT staining was performed as described by Kroschwald and collaborators (Kroschwald, Maharana et al. 2015). Cells grown to $OD_{600}$ ~0.6 were harvested and resuspended in 30 $\mu$M ThT, 10 mM Tris/EDTA buffer (pH 7) for 20 min. Cells were then washed 3 times in PBS and resuspended in LFM media for imaging. Fluorescence microscopy ($\lambda_{ex}$ of 405nm, $\lambda_{em}$ of 450/50 nm) or ($\lambda_{ex}$ of 488 nm, $\lambda_{em}$ of 525/50 nm) were performed on the Quorum platform (see Diffraction-limited fluorescence microscopy section). BY4741 cells transformed with the plasmid pRS416-GAL-Sup35NM-RFP and induced for 2 hours in 2% galactose LFM were used as positive ThT stain controls. Alternatively, we confirmed ThT *in vivo* results with ThT staining of fixed cells. Cells were fixed with 4% paraformaldehyde 2% sucrose PBS for 20 min and washed once in PBS. Cells were then permeabilized with 0.1% Triton-X phosphate buffered (pH 7.5) detergent solution, and treated with 0.001% ThT for 10 min at room temperature. ThT stained cells were washed at 3–4 times with PBS and imaged on the Quorum platform ($\lambda_{ex}$ of 488 nm, $\lambda_{em}$ of 525/50 nm) (see Diffraction-limited fluorescence microscopy section). This experiment was replicated 3 times.

**Cell treatment with water-glycerol solutions, 1,6 hexanediol and latrunculin-A**

Water and glycerol binary combinations were mixed to obtain solutions with precise osmotic pressures ranging from 1.4 MPa to 30 MPa. Cells were grown to log phase, centrifuged for 2 minutes at 3000 x g and resuspended in the different water-glycerol binary solutions in Nunc 96-well glass bottom imaging plates (Thermo Scientific; 164588). Fluorescence images of *GPD1Δ* cells, that express EGFP from a pAG416-GPD plasmid, were then captured immediately after resuspension and at 1 hour intervals with the confocal InCell6000 (see Diffraction-limited fluorescence microscopy section). Intensity thresholding of the GFP channel allowed us to segment the cells and obtain cell area values. These area values were used as a proxy for cell size at the different osmotic pressures. Measurements were taken from respective water-glycerol samples, this experiment was replicated 3 times.

We utilized 1,6-hexanediol (HD) to differentiate phase separated intracellular bodies from stable solid or fibrillar protein aggregates or to modulate the interactions among proteins that compose the endocytic condensate. HD is thought to change the solvent quality inside cells resulting in disruption of the interactions favorable to liquid-liquid phase separation. 1,2,3-hexanetriol (HT) was used as a negative control in all the HD assays. We treated cells grown to mid-log phase ($OD_{600}$ 0.6) with HD or HT from 0 to 10 % wt/v and we measured both the uptake of fluorescent probes and progression of CME protein accumulation at cortical foci by fluorescence microscopy. Sla1-GFP signal at cortical sites in the presence of HD was assessed on the Quorum platform (see Diffraction-limited fluorescence microscopy section). HD dose-response was replicated 3 times.

Inhibition of F-actin synthesis was achieved using Latrunculin A (Lat A) at concentrations determined by titrating Lat A from 0 to 200 $\mu$M. Cells were treated for 20 minutes prior to measurements of the uptake of membrane (FM6-64 dye) and formation of Abp1-mCherry fluorescence foci. Lifetime of Abp1-mCherry at cortical sites in the presence of HD was assessed on the Quorum platform (see Diffraction-limited fluorescence microscopy section) and kymographs were generated with the Metamorph image analysis software (Molecular Devices). The Lat A dose-response curve indicated that a concentration of 20 $\mu$M Lat A was sufficient to impair actin nucleation at cortical sites. We replicated this experiment 4 times. We used a concentration of 20 $\mu$M Lat A in all our experiments unless mentioned otherwise.

**Measurement of vesicle leakage of carboxyfluorescein by 1,6-hexanediol**

Vesicles containing carboxyfluorescein were produced through drying, hydration, and extrusion steps. Briefly, 0.483 mg of DOPC (Avanti Polar Lipids) and 0.007 mg of Texas Red DHPE (Life Technologies) were dried on the walls of a test tube under $N_2$ followed by 30 min of vacuum. Dried lipids were hydrated with 2 ml of 20 mM carboxyfluorescein (~20x the self-quenching concentration in Figure S2D) in 20 mM HEPES at pH ~7.39 for 2.25 h at ~60 ˚C. The resulting multilamellar vesicles were gently vortexed and then extruded in 17 passes through 100 nm pores at room temperature to make unilamellar vesicles. Unencapsulated carboxyfluorescein was removed on a Sephadex G25 size exclusion column. To produce enough sample for triplicate experiments, the procedure above was conducted twice in parallel. Yields from both runs were combined and diluted with HEPES to a final volume of 6 ml of vesicle solution. Vesicle rupture or leakage causes carboxyfluorescein concentrations to fall below the self-quenching limit, resulting in increased fluorescence. Carboxyfluorescein solutions were made in 20 mM HEPES buffer. Fluorescence was monitored with a LS-50B spectrometer (Perkin Elmer), which excited the sample at 492 nm and measured emission at 512 nm via integration for 60 s through slits widths of 15 nm. The intensity of the first data point was set to 100 (in arbitrary units). Vesicle leakage experiments were conducted in triplicate. For each run, emission at 512 nm was measured three times across a 1.0 cm path length cuvette containing 2 ml of vesicle solution to which stock solutions of 70% 1,6-hexanediol were added in order to reach 2-12% of 1,6-hexanediol by weight. Mixing was achieved by pipetting. To verify that pipetting did not disrupt vesicles, two methods were compared. For one of the three cuvettes, the solution was mixed by gently drawing 100 $\mu$l of solution into a standard 200 $\mu$l pipette tip three times. For the other two cuvettes, the solution was mixed by gently drawing 1 ml of solution into a 1 ml pipette tip that had been cut to widen the opening and decrease shear. Results from the two methods were indistinguishable. Data were normalized such that a solution with no 1,6-hexanediol has an intensity of 1.0. Control experiments were conducted in which vesicles were exposed to a high concentration (1.1%) of the surfactant Triton-X, such that all carboxyfluorescein leaked out of vesicles and was fully de-quenched.

**Direct stochastic optical reconstruction microscopy (dSTORM)**

Direct stochastic optical reconstruction data was acquired with the custom-imaging platform built by Quorum Technologies (see Diffraction-limited fluorescence microscopy section). Sample preparation for dSTORM was performed according to Ries, *et al*. with minor modifications (Ries, Kaplan et al. 2012). Cells were grown to an $OD_{600}$= 0.1 and plated on ConA coated glass bottom 35 mm round dishes for 10 minutes. Cells were then fixed with 4 % paraformaldehyde 2% sucrose PBS for 15 min. Fixation was stopped with two sequential incubations of 10 minutes in 50 mM $NH_4Cl$ PBS and cells were further permeabilized and blocked in 0.25 % Triton X-100, 5 % BSA, 0.004 % $NaN_3$ PBS for another 30 minutes. We used GFP-Booster-Atto647N nanobodies (Chromotek; code gba647n) to label Sla1-GFP at a concentration of 10 µM in 0.25 % Triton X-100, 1% BSA, 0.004 % $NaN_3$ PBS for 60 minutes. Cells were washed extensively in PBS before imaging in blinking buffer 150 mM Tris-HCl pH 8.0, 30 mM β-mercaptoethylamine (MEA), 0.5 % glucose, 0.25 mg/ml glucose oxidase and 20 µg/ml catalase. We acquired streams of 10,000 to 20,000 frames at 30 msec. exposures and we used Wave Tracer plugin (Molecular Devices) to detect and gate events with a 16-bit intensity threshold of 1,000. Measurements were taken from distinct samples to reach a count of 250 bodies and this experiment was replicated 5 times. When possible we didn't use gain on the EMCCD camera to better calculate resolution; the camera has a conversion factor of $6e^-$/count when no gain is used. Based on photon counts, we estimate an x, y-resolution of ≈10 nm with the 647 nm wavelength Atto647N fluorophore and a z-resolution of ≈50 nm with the astigmatic lens in 3D configuration calibrated on TetraSpeck beads (ThermoFisher). Center of mass for each event was calculated and we reconstructed images in Wave Tracer before further analysis in Metamorph (Molecular Devices). Sla1 structures were separated in circular and narrow elliptical shapes that correspond respectively to structure within or at the equator of cells.

**Fluorescence recovery after photobleaching**

Fluorescence recovery after photobleaching (FRAP) experiments (for recovery of Sla2-GFP and dextran-FITC) were performed on a Leica SP8 laser scanning confocal microscope (Leica). Images were acquired with a 60x oil objective and the FRAP module within the LASX confocal software (Leica). A photobleaching 488 nm laser was pulsed for 190 to 650 msec. on samples, followed by acquisition of fluorescence recovery for 1 min with time resolution between 1.5 to 5 frames per seconds. GFP or FITC signal recovery was measured within either a segmented Sla1-mCherry or Syp1-mCherry region of interest to ensure that FRAP was acquired within the endocytic condensate. We also measured recovery in neighbor cytosol regions to assess recovery of dextran-FITC outside condensates. Analysis of the images intensity (I) fluctuations and segmentation of regions of interest (ROI) were performed on the LASX imaging software (Leica). Recovery measurements were taken from distinct samples. We replicated the dextran-FITC experiment 6 times and the Sla2-GFP recovery 4 times. We analyzed the data as follows:

We first applied a double normalization on bleached ROI1:

$$I(t)_{dbl\,norm} = \left( \frac{\frac{1}{n_{pre}} \cdot \sum_{t=1}^{n_{pre}} I(t)_{ROI\,2'}}{I(t)_{ROI\,2'}} \right) \cdot \left( \frac{I(t)_{ROI\,1'}}{\frac{1}{n_{pre}} \cdot \sum_{t=1}^{n_{pre}} I(t)_{ROI\,1'}} \right); \qquad (1.1)$$

, where ROI2 is a non-bleached region and $n_{pre}$ is the number of pre-bleached images. We then used (1.2) results to perform a full-scale normalization:

$$I(t)_{full\,norm} = \frac{I(t)_{dbl\,norm} - I(t_{post})_{dbl\,norm}}{I(t_{post})_{dbl\,norm}} \qquad (1.2)$$

, where the first post bleach data time points are given a value of zero. We finally fitted each normalized trace with a non-least square function to best fit the single term equation:

$$f(I) = I_0 - a \cdot e^{-\omega t}\,; \qquad (1.3)$$

from which we extracted mobile fractions and half recovery times for the Sla2-GFP and dextran-FITC samples. Analyses were performed using subroutines of the R package.

**DHFR PCA procedure**

We performed the DHFR PCA procedure has previously described (Tarassov, Messier et al. 2008) with minor modifications for a selected subarray of 30 potential Sla1 interactors from Biogrid. *MAT***a** strains harbouring either *Sla1-DHFR-F[1,2]-nat1* or *Sla1-ΔPLD-DHFR-F[1,2]-nat1*fusions were grown in liquid YPD supplemented with 100 $\mu$g/mL nourseothricin in a square v-bottom 96-well array block (3960; Corning). Each *MAT***a** strain was distributed in 3 rows by 10 columns arrangement to obtain 6 rows by 10 columns array on a single 96-well block. Another block array (6 rows by 10 columns arrangement) of 30 different *MAT***α** strains with *ORF.X-DHFR-*

*F[3]-hph* fusions was cultured in liquid YPD with 250 $\mu$g/mL hygromycin B. The strains were grown in liquid array for 24 hours at 30°C. Mating procedure was performed in a 96-well liquid array (6 rows by 10 columns arrangement) for 24 hours at 30°C. Each *MAT***a** and *MAT***α** strains were combined one-to-one in fresh liquid YDP without selection for mating, so that each resulting diploid strain had respective genes tagged with either of the DHFR fragments. The array was then transferred in quadruplicate in a 1,536-format using a robotically manipulated 384-pin tool (0.787 mm flat round-shaped pins, custom AFIX384FP3 BMP Multimek FP3N, V&P Scientific Inc., San Diego, CA) to synthetic complete agar plates (omnitrays, Nunc™) without lysine and without methionine and with 100 $\mu$g/mL nourseothricin and 250 $\mu$g/mL hygromycin B and were incubated for 48 h at 30°C to select for diploid cells. This step was repeated to further select for diploid cells using a robotically manipulated 1,536-pin tool (0.457 mm flat round-shaped pins, custom AFIX1536FP1 BMP Multimek FP1N, V&P Scientific Inc., San Diego, CA) to transfer the array on fresh plates. For selective growth, the strains were printed in 1,536 format on SC agar medium without adenine (pH 4.8) containing 4 % (w/v) Noble agar, 2 % (w/v) glucose, 1.74 g/L YNB without ammonium sulfate, and methotrexate (200 $\mu$g/mL) to select for DHFR reconstitution with either *Sla1-DHFR-F[1,2]-nat1* or *Sla1-ΔPLD-DHFR-F[1,2]-nat1* baits. We repeated the methotrexate selection on 3 distinct replicate plates. Printed methotrexate plates were incubated at 30°C throughout the imaging process and individual plates were photographed each day for 7 days with a 4 megapixel Canon digital camera (Powershot A520). Quantification of colony growth was achieved as described in Stynen *et al.* 2018 (preprint available on bioRxiv) with a custom-made macro on imaging software Fiji (v1.45b) to extract the integrated density of each colony. Briefly, for each colony, a 17-by-17 pixel area was screened around an initial central position to find the central intensity gravity point. From this central point, we scanned for the first pixels with intensity values that equal background intensity in the left, right, bottom and up directions and used these cardinal coordinates to determine a new central pixel. We iterated this process to get the borders of the colony in all four directions and a final central pixel. An oval was created and analyzed for integrated density, which is the mean density times the area. Further analyses were performed using a custom-made script in R (v3.2.3). The integrated densities calculated in the previous section were $\log_2$ transformed.

**Phase separation and partitioning of fluorescent proteins in cleared cell lysates**

Yeast cells that respectively express pellets were harvested and disintegrated by Y-PER reagent. Pellets were resuspended in the Y-PER reagent solution and incubated at room temperature with shaking for 20min. The homogenate was centrifuged at 20 000×g for 30min and the pellet was discarded (see Thermo scientific YPER protein extraction instructions). Homogeneous cleared cell lysates were titrated with PEG from 0 to 10% and monitored by confocal microscopy (see Quorum platform) for appearance of a condense droplet phase. Fluorescent signal from respective protein was acquired along the brightfield channel with a HCX PL APO 63x /1.47 NA oil corrected TIRF objective (Leica) and 50 $\mu$m pinhole spinning disk on our Quorum Diskovery platform (see Diffraction-limited fluorescence microscopy section).

**Expression of chimeric Epsins, mammalian Epsin homologs and non-cognate PLDs**

Sequence encoded information, shown by the meta-predictor PONDR-FIT(Xue, Dunbrack et al. 2010), suggest that yeast proteins Ent1 and Ent2 and their mammalian homologs EPN1 and EPN3 are in fact intrinsically disordered proteins (IDPs) with defined disordered (*low complexity*) domains that are conserved despite that the corresponding sequences not being conserved. For Ent1 and Ent2 the predicted disordered domains match with previous predictions for prion domains (PLDs) (Alberti, Halfmann et al. 2009).

We expressed homologous human proteins EPN1 and EPN3 in yeast. Human ORFs from the CCSB Human ORFeome library from Dana-Farber Cancer Institute (Dharmacon) were sub-cloned into the pAG yeast expression vectors with a fluorescent tag. Target ORF were cloned in the yeast gateway expression vector pAG416GDP-ccdB-EGFP (Addgene) from their respective pDONR entry plasmids. Sequence variants of Sup35 NM (Sup35 NM, Sup35 NM(N) and Sup35 NM(Q)) and Sla1 PLD were also cloned in the yeast gateway expression vectors pAG416GDP-ccdB-EGFP (Addgene) from their respective pDONR entry plasmids (generous gift from R Halfmann) and tested for colocalization with Sla1 condensates. Finally, ENTH domains of Ent1/ and EPN1/3 were cloned between the XbaI and SpeI sites of the pAG416GDP-*ORF.X*-EGFP vectors to create chimeric Epsin proteins with either the Sla1 PLD, Sup35 NM, Sup35 NM(N) or Sup35 NM(Q) moiety. We tested colocalization of each construct with the endocytic condensate (BY4741 SLA1-mCherry::*hph* cells) and functional complementation. Images were acquired with a HCX PL APO 63x /1.47 NA oil corrected TIRF objective (Leica) and 50 $\mu$m pinhole spinning disk on our Quorum Diskovery platform (see Diffraction-limited fluorescence microscopy section).

**Centroid tracking of Sla1 foci**

We measured the mean square displacement (MSD) of single Sla1-YFP fluorescent foci within a confocal volume on the Quorum platform (see Diffraction-limited fluorescence microscopy section). Images were acquired at 20 fps for 30 seconds with a 50 $\mu$m pinhole spinning disk and we performed particle centroid tracking with the Wave Tracer plugin in the Metamorph software (Molecular Devices). We included only foci that remained in the confocal volume throughout the acquisition and which showed decreasing fluorescence intensity as a function of MSD (Figure S4I). We next determined the linear displacement of the Sla1 foci towards the cell interior with the particle coordinates plotted as a function of elapsed time (Figure S4J). Measurements were taken from multiple cells in a single sample and this experiment was replicated 3 times.

**Quantification of membrane in nascent vesicles under HD titration**

We quantified the amount of membrane in single nascent CME vesicles by fluorescence emitted from FM6-64 labelled membrane. Overnight *GPD1*Δ Sla1-YFP cell cultures were diluted 1:40 in fresh LFM with HD concentrations from 0 to 5%. Cells were first incubated in 20 $\mu$M Lat A and then in the HD solutions for 5 minutes and then labelled with 5 μM FM4-64 for another 5 minutes before direct fluorescence image acquisition on the Quorum platform (see Diffraction-limited fluorescence microscopy section). Single vesicles where segmented with an intensity threshold in both Venus YFP and FM4-64 channels to quantify the membrane fluorescence that co-localizes with Sla1 signal. Measurements were taken from multiple cells in a single sample and this experiment was replicated 5 times. Average intensity measurements per nascent vesicle were normalized to values between 0 and 1 for the whole HD treatment concentration range. As a reference point to compare with membrane invagination predictions, we extracted mean and standard deviation values from data for HD concentration below 2 %; mean and standard deviation were also determined for each HD concentration.

**Effect of 1,6-hexanediol (HD) titration on condensate stability**

Formation of a endocytic condensate results in an interface between the condensate and the cytosol and we can treat the condensate and the dispersed cytosol as two phases defined by an interface between them. In a mean field description, the cohesive interactions that drive the formation of the condensate derive from the balance of interactions amongst the condensate and cytosolic components. This will determine the stability of the condensate. In addition, the interface between the condensate and cytosol will be governed by an interfacial tension. A simple adaptation of the Flory-Huggins model for binary mixtures can be used to quantify the interfacial tension (Dill and Bromberg 2011).

The model is as follows: We shall define two condensed phases *viz*., the condensate phase (D) and the cytosolic phase (C). The interfacial tension $\gamma_{DC}$ defines the free energy penalty associated with increasing the interfacial area between the two phases. If $\gamma_{DC} > 0$, then the interfacial area will be minimized, thus resulting in spherical condensates. From the vantage point of the condensate, reducing the interfacial tension decreases the number of condensate components that are "sacrificed" to be at the interface and thus lose favorable intra-condensate interactions.

If the total free energy of the two bulk phases D and C and the interface between the phases is $F$, then the interfacial tension associated with changing the interfacial area $A$ is defined as:

$$\gamma_{\mathrm{DC}} = \left(\frac{\partial F}{\partial A}\right)_{n_D, n_C, T} ; \qquad (2.1)$$

For simplicity, we shall use a mean-field model with the two phases defined on a lattice with coordination number $z$. The molecules of D and C will be considered to be of similar size and the translational entropy will be set to zero. If $a$ is the area per molecular unit that is exposed to the interface, then equation (2.1) becomes:

$$\gamma_{\mathrm{DC}} = \left(\frac{\partial F}{\partial A}\right)_{n_D, n_C, T} = \left(\frac{\partial U}{\partial A}\right)_{n_D, n_C, T} = \frac{1}{a}\left(w_{\mathrm{DC}} - \frac{w_{\mathrm{DD}} + w_{\mathrm{CC}}}{2}\right); \qquad (2.2)$$

Here, the $w$ terms are the effective mean-field energies associated with interactions between components of the condensate ($w_{\text{DD}}$), the cytosol ($w_{\text{CC}}$) and the components of the condensate and the cytosol ($w_{\text{DC}}$). These energies are in units of $k_BT$ and the convention is that the energies are negative if they are favorable and positive if they are unfavorable. Accordingly, it follows that:

$$\begin{aligned}\gamma_{\text{DC}} > 0 \text{ if } \left|w_{\text{DC}}\right| < \left|\frac{w_{\text{DD}} + w_{CC}}{2}\right| \\ \gamma_{\text{DC}} = 0 \text{ if } \left|w_{\text{DC}}\right| = \left|\frac{w_{\text{DD}} + w_{CC}}{2}\right|; \\ \gamma_{\text{DC}} < 0 \text{ if } \left|w_{\text{DC}}\right| > \left|\frac{w_{\text{DD}} + w_{CC}}{2}\right|\end{aligned} \tag{2.3}$$

Importantly, equation (2.2) can be rewritten in terms of the Flory-Huggins interaction coefficient using the relationship:

$$\chi_{\text{DC}} = \frac{z}{k_BT}\left[w_{\text{DC}} - \left(\frac{w_{\text{DD}} + w_{\text{CC}}}{2}\right)\right] = \left(\frac{c_E}{k_BT}\right); \tag{2.4}$$

Here, $c_E$ is cohesive energy that holds the condensate together and represents the balance of condensate-cytosol, intra-condensate and intra-cytosol interactions. Accordingly,

$$\begin{aligned}\gamma_{\text{DC}} = \frac{1}{a}\left(w_{\text{DC}} - \frac{w_{\text{DD}} + w_{\text{CC}}}{2}\right) = \left(\frac{k_BT}{za}\right)\chi_{\text{DC}}; \\ za\gamma_{\text{DC}} = k_BT\chi_{\text{DC}}\end{aligned} \tag{2.5}$$

Alternatively,

$$\chi_{\text{DC}} = \left(\frac{za}{k_BT}\right)\gamma_{\text{DC}} = \left(\frac{c_E}{k_BT}\right); \tag{2.6}$$

$$za = \left(\frac{c_E}{\gamma_{\text{DC}}}\right)_{\text{HD=0\%}}; \tag{2.7}$$

Note that the values of $z$ and $a$ are fixed by the lattice and components of the condensate. Through measurements combined with the Young-Laplace theory, we have estimates of the interfacial tension and $c_E$ in the absence of HD – as shown in equation (2.7). These estimates can be used as shown in equation (2.7) to estimate the value of $za$. Since this value of $za$ is independent of HD concentration, one can fix $za$ and use the estimate of cohesive energy $c_E$ at different HD concentrations to estimate the change in interfacial tension as a function of HD concentration using equation (2.8) below:

$$\gamma_{\text{DC}}(\%\text{HD}) = \left[\frac{c_E(\%\text{HD})}{za}\right]; \tag{2.8}$$

**Polystyrene beads and Dextran-FITC osmoporation**

To incorporate dextran-FITC of different chain length inside haploid yeast *GPD1*Δ cells, we used an osmoporation technique similar to that described by da Silva Pedrini *et al*. (da Silva Pedrini, Dupont et al. 2014). Cells treated with Lat A were centrifuged for 2 minutes at 3000 x g and resuspended in a water-glycerol binary solution at 1.4 MPa for 30 minutes, then in a 30 MPa solution for 1 hour. Osmoporation of dextran-FITC is performed after these steps, by centrifuging cells at 3000 x g and resuspending the pellet in the 1.4 MPa water-glycerol solution with the dextran-FITC at 10 mg/ml for 1h. After this incubation period, cells were put on ice and washed 3 times with cold PBS with 20 $\mu$M Lat A. Cells were preserved on ice until they were plated on ConA treated 35 mm imaging dishes for 10 minutes and imaged on either the Quorum or Leica platforms (see Diffraction-limited fluorescence microscopy and Fluorescence recovery after photobleaching sections respectively).

We also incorporated 200 nm orange (540/560) FluoSpheres carboxylate-modified polystyrene beads (ThermoFisher) with this technique. Beads where either coated with 10 mg/ml BSA prio to incorporation into cells. After osmoporation, cells were preserved on ice until they were plated on ConA treated coverslips and mounted on glass imaging slides for subsequent image acquisition and optical tweezers experiments.

**microNS-GFP micro-rheology**

To determine the effect of *osmoporation* on the cell rheological properties, we measured displacement of expressed viral capsid microNS particles labeled with GFP in both normal and osmoporated cells (Figure S4A-F). Cells were transformed with the microNS-GFP pRS expression plasmid, a generous gift of S. Alberti at Max Planck Institute of Molecular Cell Biology and Genetics (MPI-CBG). microNS movement was recorded on the Quorum platform (see Diffraction-limited fluorescence microscopy section) within a 2 microns thick Z-stack of 5 confocal planes (50 $\mu$m pinholes) acquired at 5 frames per second (fps). Measurements were taken from multiple cells in a single sample and this experiment was replicated 3 times. Image analysis was performed with Metamorph and Wave Tracer plugin (Molecular Devices), to track particles displacement with centroid localization on maximum intensity projections. We further filtered the mean square displacement (MSD) data for particles that are confined and showed that they have MSD similar to the technical noise of our apparatus setup.

**Optical tweezers measurements and calibration**

Dynamic mechanical analysis of yeast cytoplasm was performed with a custom optical tweezers (OT) platform. Our OT system is an inverted microscope (Nikon Ti-E) equipped with a CFI APO SR TIRF 100x / 1.49 NA oil immersion objective (Nikon), a 1,064 nm Nd:$YVO_4$ 10 W infrared laser (IPG Photonics), an X-cite lamp source (Excelitas) and a nano-positioning stage (Mad City Labs). Oscillation of the tweezers on the specimen plane from 0.1 Hz to 2000 Hz was achieved with an acousto-optic deflector (AOD, AA Optoelectronics) coupled to a digital frequency synthesizer that we controlled with in house Labview (National Instruments) routines. Light transmitted through the specimen was collected with a condenser lens and reflected onto a position-sensitive detector (PSD) (Thorlabs, PDP90A) to perform back focal plane interferometry. Before acquisitions we adjusted the microscope for Köhler illumination and ensured that all the optics were conjugate to the respective specimen plane or back focal plane. At each frequency of excitation we recorded the signals (120,000 samples at 1000 Hz to 2,000,000 samples at 0.1 Hz at 20 kHz) and performed Fourier analysis. Measurement time for each frequency sweep was about 15 minutes on distinct samples and this experiment was replicated 2 times. For each sample, we covered the frequency domain from high-to-low frequencies, then we repeated the procedure from low-to-high to ensure consistent frequency response with prolonged laser exposure.

Calibration of the optical tweezers measures was performed as previously described with minor modifications (Fischer, Richardson et al. 2010, Hendricks, Holzbaur et al. 2012). Data analysis was conducted with in-house Matlab code. Data quality was first confirmed by assessment of the sinusoidal shape of the response to the applied stress. Traces with a coherence of 0.95 or greater were included in the analysis. We averaged 17 traces in distinct cellular locations and determined their trap stiffness $k_{trap}$ (mean ± standard error; 8.0 x $10^{-5}$ ± 2.7 x $10^{-5}$ N•m$^{-1}$), photodiode sensitivity factor $\beta$ (mean ± standard error 10.7 x $10^3$ ± 2.3 x $10^3$ nm•V$^{-1}$) and frequency-dependent viscoelastic moduli G’ (storage) and G’’ (loss) (Figure 4).

To account for the relaxation dynamics observed, we performed a fit on the forced response to sinusoidal oscillations and the power spectra of the spontaneous fluctuations of the bead to a model of a transiently crosslinked network of flexible polymers (Lieleg, Schmoller et al. 2009) .

$$G' = G_0 - a \cdot \frac{N k_{off}}{\frac{k_{off}^2}{4\pi^2} + f^2} + b \cdot \left(\frac{f}{f_0}\right)^{\alpha} ; \qquad (2.9)$$

$$G'' = c \cdot \frac{N f}{\frac{k_{off}^2}{4\pi^2} + f^2} + d \cdot \left(\frac{f}{f_0}\right)^{\alpha} ; \qquad (2.10)$$

From these fits, we extracted the average parameters ($G_0$; 16.53), ($f_0$; 37.44), number of crosslinks (N; 3 x $10^{14}$), crosslink off-rate ($k_{off}$; 8.28), power law ($\alpha$; 0.98) and constants ratios (a/b; 1.06 x $10^{-14}$/2.19), (c/d; 1.33 x $10^{-14}$/0.72). Deviation of $\alpha$ above a value 0.75 indicates that retraction or extension of entangled protein filaments ends could contribute to the relaxation mechanism (Koenderink, Atakhorrami et al. 2006).

**Dimensions and geometry of membrane contour and endocytic condensates**

From published data, we considered that the optimal membrane U-shaped geometry before vesicle excision is about 70 nm high and 60 nm wide (Idrissi, Blasco et al. 2012). To approximate the area and volume, the U-shape was decomposed into a hemispherical cap of 30 nm radius over a cylinder of 30 nm radius and 40 nm high. We then calculated, with these dimensions, an invaginated membrane area of 1.32 x $10^{-14}$ $m^2$ and volume of 1.70 x $10^{-22}$ $m^3$.

We also defined for the membrane bending energy, the invagination height profile as a function of position around the invagination peak and middle of the condensate;

$$h(x,y) = h_0 \cdot \exp\left(\frac{-(x^2+y^2)}{2R_0^2}\right); \tag{3.1}$$

where $h_0$ is the invagination depth and $R_0$ is the radius of invagination.

To calculate the area and volume occupied by the endocytic condensate and thus the amount of displaced cytosol material, we used our dSTORM measurements. The hemispherical endocytic condensate volume was calculated to be 2.43 x $10^{-21}$ $m^3$ for a radius of 105 nm. The same hemispheric condensate has an area of 6.93 x $10^{-14}$ $m^2$. These dimensions agree with the size of ribosome exclusion zones observed surrounding invaginated clathrin patches observed by EM (Kukulski, Schorb et al. 2012).

**Calculation of the energies that favor and counteract membrane invagination**

We calculated the membrane bending energy with the standard Helfrich model for the membrane profile obtained by equation (3.1) and considered a membrane bending modulus $\kappa_m$ of 12.5•$K_B$T (Helfrich 1973, Harmandaris and Deserno 2006, Carlsson and Bayly 2014):

$$U_{em} = 11\sqrt{\left(\frac{\pi^3}{32}\right)}\left(\frac{h_0}{R_0}\right)^2 \kappa_m; \tag{3.2}$$

We determined the energy required to generate a 70 nm deep invagination to be 3 x $10^{-18}$ J.

We considered that the viscoelastic cytosol behaves as a Kelvin-Voigt material and that the total stress is equal to the sum of the elastic and viscous stresses, such that:

$$\sigma = \varepsilon E_i + \dot{\varepsilon}\eta; \tag{3.3}$$

, where η is the cytosol viscosity at a specific frequency $f_x$ obtained with:

$$\eta = \frac{G''}{2\pi f_x};\qquad (3.4)$$

For stringency, we used a Young's (or elastic) modulus (E) for the cytosol of 45 Pa (see Eq. 3.6 below) and calculated the deformation with the condensate radius ($1.18 \times 10^{-7}$ m) over the cell radius ($2 \times 10^{-6}$ m). We obtained an elastic stress of 3 Pa.

We next determined η at 0.5 Hz to be 0.35 Pa•s. and the deformation rate of 0.004 $s^{-1}$, for a condensate velocity $v$ of $7.4 \times 10^{-9}$ m•$s^{-1}$ (Figure S4I,J).

$$\dot{\varepsilon} = \frac{v}{6\pi R_{drop}};\qquad (3.5)$$

, which gives a negligible viscous stress of 0.0014 Pa. With a volume of $2.42 \times 10^{-21}$ $m^3$, the total stress of 3.0014 Pa corresponds to an energy cost of $7.26 \times 10^{-21}$ J to displace the viscoelastic cytosol.

**Endocytic condensate material properties and contact angle**

We determined the values of the elastic modulus of the cytosol based on the observation that compressible biological materials have a Poisson's ratio (ν), which relates all material moduli to each other, of 0.45 (or between 0.3 and 0.5) (Zhang, Soman et al. 2013). We then used the relationship between shear modulus (G) and Young's (or elastic) modulus (E):

$$E_i = 2(1 + \nu_i)G_i;\qquad (3.6)$$

to calculate an approximate elastic modulus of 45 Pa for the cytosol from the G' of 15 Pa at 1 Hz. This strain rate was used to consider a G' near the elastic plateau of the material response. With the volume of displaced cytosol and this modulus value, we could determine the mechanical stress imposed on the cytosol to be ~3 Pa (see Eq. 3.3-3.5). This corresponds to a compression force (F) of ≈$1.4 \times 10^{-13}$ N. We then isolated the equivalent modulus (E`) from Hertz elastic contact equation

$$F = \frac{4}{3}E'_{ij}R'^{\frac{1}{2}}_{ij}\delta^{\frac{3}{2}}_{ij};\qquad (3.7)$$

into

$$E'_{ij} = \frac{4F}{3R'^{\frac{1}{2}}_{ij}\delta^{\frac{3}{2}}_{ij}};\qquad (3.8)$$

, where $\delta_{ij}$ is the interface indentation (or invagination depth), to estimate the respective elastic moduli of the cytosol and the condensate, again with both materials having a Poisson's ratio of 0.45. The equivalent modulus is determined by:

$$\frac{1}{E'_{ij}} = \frac{(1+\nu_i)^2}{E_i} + \frac{\left(1+\nu_j\right)^2}{E_j}; \tag{3.9}$$

where $E_i$ and $E_j$ are the respective elastic moduli of the objects in contact and

$$\frac{1}{R'_{ij}} = \frac{1}{R_i} + \frac{1}{R_j}; \tag{3.10}$$

for contact between two spheres of radii $R_i$ and $R_j$. These relationships gave us a endocytic condensate elastic modulus of 59 Pa.

Given that gravity is negligible at the scales we are measuring, the condensate can be treated as a sectional arc of a sphere or hemispherical cap alone (Figure S1G) and we can estimate the condensate contact angle;

$$\theta = 2 \cdot \tan^{-1}\left(\frac{h}{d}\right) \tag{3.11}$$

, where d is condensate radius and h is condensate height. Based on our 2D dSTORM measures we obtained a contact angle of about 97°. Note that at the hundred nanometer scale, condensate wetting geometries are also affected by evaporation and line tension at the three-phase contact line.

**Theoretical model based on elastic and adhesive contact mechanics**

To explain how phase separation of disordered proteins into a 100 nm-scale viscoelastic body can invaginate the membrane (in the absence of turgor, F-actin polymerization and steric effects), we propose a model in which elastic and adhesive contact mechanics can generate enough energy (estimated from membrane bending to be $\approx 3 \times 10^{-18}$ J) to drive local invagination of the membrane.

We explored the idea that when endocytic condensates are nucleated between the membrane and cytoplasm, new interfaces are created and the free energy available on the condensate interfaces can produce work to deform both surrounding substrates (membrane and cytosol). We thus hypothesize that free energy is released upon phase separation and adhesion of the endocytic condensate to neighbor structures, which is converted into mechanical work. We describe below the details of the model.

We first propose a general model where the energy penalties to create interfaces around the condensate and deform the cytosol and the membrane follow a super-linear growth as a function of invagination depths $\delta$ (of both cytosol and membrane), whereas the free energy released by condensate phase separation is linear. We expressed the relationship between these two general energy terms and the total energy $U$ with the power-law function:

$$U \sim \phi \cdot \delta^{1+\varepsilon} - \psi \cdot \delta; \tag{4.1}$$

, where $\phi$ is the energy penalty term, $\psi$ is the available work and the exponent $\varepsilon > 0$. When $\delta$ is isolated from the partial derivative of equation (4.0), such as:

$$\frac{\partial U}{\partial \delta} = \phi(1+\varepsilon) \cdot \delta^{\varepsilon} - \psi;$$

$$\delta^{*} = \left(\frac{\psi}{\phi(1+\varepsilon)}\right)^{\frac{1}{\varepsilon}}; \tag{4.2}$$

, we observe that our model predicts that $\delta$ scales with $\phi$ and $\psi$:

$$\delta^{*} \sim \frac{\psi}{\phi}; \tag{4.3}$$

We provide below a proof for this model, where $\phi$ is decomposed into individual elastic, viscous friction and surface stress terms and $\psi$ is fragmented in the work of adhesion from the respective condensates interfaces. We also describe the quantities contained within these individual terms; either directly measured, calculated or estimated.

Parallel to the membrane plane, surface energies formed by the condensate (D), cytosol (C) and membrane (m) come to equilibrium as described by Young's equation:

$$\gamma_{cm} = \gamma_{dm} + \gamma_{dc} \cdot \cos\theta; \tag{4.4}$$

With a contact angle of about 97°, the cytosol-membrane surface tension ($\gamma_{cm}$) is predicted to be approximately equal to the condensate-membrane surface tension ($\gamma_{dm}$). We determined the values of $\gamma_{dc}$ and $\gamma_{dm}$ next.

We used an apparent Young's modulus ($E_{cell}$) of 1 kPa, determined from atomic force microscopy on haploid yeast spheroblast (Munder, Midtvedt et al. 2016). We considered that this modulus represents the bulk material properties of both membrane and cytosol when deformed towards the cell interior, as for a CME-driven invagination. We then calculated the overall mechanical stress in the system with Hooke's law:

$$\sigma = \varepsilon E_i; \tag{4.5}$$

to create the U-shape membrane geometry observed by EM. The deformation ε of the membrane was decomposed to represent stretching of an elastic band, where deformation equals length difference over original length. For our U-shaped deformation the linear contour was established at 174.2 nm, whereas the same membrane line measured ≈ 80 nm before invagination. This gives us a deformation of 1.18 (dimensionless) and a stress of 1178 Pa.

This mechanical stress corresponds to the pressure difference ΔP experienced by the cytoplasm and the membrane due to the presence of the endocytic condensate. This pressure difference arises from adhesive, hydrostatic and elastic stress energies. We then used the Young-Laplace equation:

$$\Delta P = 2\gamma_{dc} \cdot H \; ; \tag{4.6}$$

, where H is the mean curvature of the interface and is equal to $1/R_d$, to calculate $\gamma_{dc}$ at the cytosol interface based on our observation that the condensate is circular in shape. Equation (4.6) determines the pressure difference across the condensate-cytosol curved interface as a function of surface energy, and gives a condensate/cytosol interfacial tension $\gamma_{dc}$ of $7 \times 10^{-5}$ N•m$^{-1}$.

Based on Young's equation (4.4) and the endocytic condensate-membrane contact angle θ value, the relationships between interfacial tensions is:

$$\gamma_{cm} = \gamma_{dm} + \gamma_{dc} \cdot \cos\theta$$
$$\gamma_{cm} = \gamma_{dm} - (8 \cdot 10^{-6}); \tag{4.7}$$

and

$$\gamma_{dc} > \gamma_{dm}; \tag{4.8}$$

where the condensate-membrane surface tension $\gamma_{dm}$ can range between about 1.8 to $7 \times 10^{-5}$ N•m$^{-1}$. The $\gamma_{dm}$ limits also arise from equation (4.9) below.

The interfacial tension γ of a fluid condensate is inversely proportional to the square of the length ξ of its discrete elements (i.e. the individual biopolymers), we used this relationship to estimate maximal and minimal γ values:

$$\gamma = \frac{k_B T}{\xi^2}; \tag{4.9}$$

, where $k_B$ is the Boltzmann constant and T is temperature in kelvin (Brangwynne, Mitchison et al. 2011). The interfacial tension values for protein based biological condensates (in which protein radii range from 1 to 10 nm) should be on the order of $1 \times 10^{-4}$ N•m$^{-1}$ and below.

To directly estimate the work of adhesion at the endocytic condensate interfaces, we used the Young-Dupré equation:

$$W_{dm} = \gamma_{dc} \cdot (1 + \cos\theta); \quad (4.10)$$

that implies a direct relationship between work of adhesion of an interface ($W_{ij}$), the condensate contact angle and the surface tension $\gamma_{dc}$. We could determine the work of adhesion at the membrane interface $W_{dm}$ to be 6 x $10^{-5}$ N•m$^{-1}$.

We also combined the Young's equation (4.1) and the Dupré relationship:

$$W_{dc} = \gamma_{dm} + \gamma_{cm} - \gamma_{dc}; \quad (4.11)$$

to express the work of adhesion at the endocytic condensate-cytosol interface $W_{dc}$:

$$W_{dc} = 2\gamma_{dm} + \gamma_{dc} \cdot (\cos\theta - 1); \quad (4.12)$$

in terms of interfacial tension at endocytic condensate interfaces with both membrane and cytosol. We could thus estimate the maximal $W_{dc}$ value to be 6 x $10^{-5}$ N•m$^{-1}$, if $\gamma_{dm} \approx \gamma_{dc}$.

We determined in parallel whether adhesion dominates the mechanical potential of endocytic condensates, as opposed to capillary effects. Under so called elasto-capillary action, condensate interfacial tension (Y) can deform an elastic sheet as a function of either condensate radius (R) or the thickness (h) and elastic modulus (E) of the slender material in contact with the condensate; in our case, the plasma membrane (Roman and Bico 2010). We can considered that the phospholipid bilayer membrane is about 10 nm thick and the cell has a bulk elastic modulus E of 1 kPa (Munder, Midtvedt et al. 2016). With a surface tension $\gamma_{dc}$ of 7 x $10^{-5}$ N•m$^{-1}$, we determined that R is larger and h is about equal to the elasto-capillary length:

$$\frac{\Upsilon}{E}; \quad (4.13)$$

, and the membrane sheet should remain undeformed at the 3-line contact point by interfacial tension alone (Figure 6A). This prediction is also consistent with EM data where the membrane doesn't bulge out under the Laplace pressure within condensates on cortical sites (Idrissi, Grotsch et al. 2008). In this scenario our system should obey the Young-Dupré equation and deformation can only come from work of adhesion.

Johnson-Kendall-Roberts (JKR) theory describes how non-flat surfaces stick together and conform to one another to minimize their interfacial energy (Johnson 1971). When they adhere to one another, soft and compliant materials such as the membrane and cytoplasm are subject to a deformation limited by elastic strain. Style, *et al.* adapted the JKR theory of contact mechanics to describe the contact surface geometry between a microscopic rigid particle and a soft substrate (Style, Hyland et al. 2013); we followed a similar approach to estimate model 4.1 parameters and test our hypothesis.

If we consider the two condensate interfaces where deformation occurs, both membrane and cytoplasm, the energy penalty to create these curved surfaces (or interfaces) are equal to the sum of elastic, viscous and surface energies.

To build a complete energy model, the elastic energy penalties were determined with the JKR theory. Since the geometry of the contact surface corresponds to the endocytic condensate geometry itself, we calculated the elastic penalties $U_{ely}$ to deform both interfaces as a function of invagination depth $\delta_{ij}$:

$$U_{e|y}\left(\delta_{ij}, E'_{ij}, R'_{ij}\right) = cE'_{ij}R'^{\frac{1}{2}}_{ij} \cdot \delta_{ij}^{\frac{5}{2}}; \tag{4.14}$$

where $E'_{ij}$ and $R'_{ij}$ are calculated with (3.5) and (3.6) respectively and c is a constant.

$$c = \frac{8}{15} \cdot \sqrt{3}\,; \tag{4.15}$$

We incorporated a correction for the membrane elastic penalty to compensate for the reduced JKR accuracy at the hundred nanometer scale and for soft materials (Style, Hyland et al. 2013) by addition of the Helfrich Hamiltonian of membrane bending at individual $\delta_{ij}$ values. We substituted $h_0$ in equation (3.2.0) for $\delta_{ij}$ and used a fixed radius of invagination $R_{ij}$ to get the relationship:

$$U_{em.corr}\left(\delta_{ij}\right) = 11\sqrt{\left(\frac{\pi^3}{32}\right)}\left(\frac{\delta_{ij}}{R_{ij}}\right)^2 \kappa_m; \tag{4.16}$$

We calculated a corrected energy cost of ≈1 x $10^{-18}$ J to deform the membrane (Table S3). The cytoplasm elastic energy barrier did not require correction and we considered no elastic nor viscous condensate deformation in our model.

JKR theory is also less accurate for very soft materials and small particles because the model neglects surface stresses (Style, Hyland et al. 2013). Inclusion of energy penalties to increase surface length at the interfaces was proposed by Style *et al.* to compensate for this reduced accuracy (Style, Hyland et al. 2013). We incorporated the surface penalties ($U_{ily}$) for the formation of the new interfaces into our model with the function;

$$U_{i|y}(\delta_{ij}, \gamma_{ij}) = \pi\gamma_{ij} \cdot {\delta_{ij}}^2 \quad (4.17)$$

, where the respective interfacial tensions $\gamma_{dc}$ and $\gamma_{dm}$ determine the energy cost to increase the interfacial areas. The energy to form the condensate/cytosol interface is ≈1 x $10^{-18}$ J and is equivalent to the combined elastic penalties. These energy values confirm that at the hundreds of nanometers scale, surface stress can dominate (or equate) elasticity in material responses to deformation. With this approach, we propose a more comprehensive model of the energy required to deform the membrane on cortical sites, that also takes into account creation of new surfaces and deformation of the cytosol.

We also incorporated an additional energy penalties $U_{v|y}$ to displace the viscous cytosol with the equation:

$$U_{v|y}(\delta_{ij}, R'_{ij}, \eta, \dot{x}) = \eta\dot{x}\delta_{ij} \cdot 6\pi R'_{ij}; \quad (4.18)$$

, where the displacement rate is the condensate maximum velocity of 7.4 x $10^{-9}$ m•$s^{-1}$ (Figure S4I,J).

If the conversion of the energy released by adhesive contact into mechanical energy is above the total energy barrier, the condensate should drive membrane invagination. The extent of membrane invagination will be limited by the free energy available. For the purpose of our model, we calculated the energy of adhesion ($U_{a|y}$) with the JKR term:

$$U_{a|y}(\delta_{ij}, W_{ij}, R'_{ij}) = 2\pi W_{ij} R'_{ij} \cdot \delta_{ij}\ ; \quad (4.19)$$

where $W_{ij}$ is the work of adhesion at each interface, as determined by equations (4.10) and (4.12), respectively. The work of adhesion refers to the energy released in the wetting process of the endocytic condensate on the membrane, it equals the work needed to separate the two adjacent phases and is given by the Dupré equation (4.11).

We then integrated the elastic (4.14, 4.16), interfacial (4.17), viscous (4.18) and adhesion (4.19) terms into a complete energy equation for i,j interfaces, where:

$$i = \{droplet\};$$
$$j = \{membrane, cortex\}; \quad (4.20)$$

into

$$U_{total}(\delta_{dm}, \delta_{dc}, \gamma_{dm}, \gamma_{dc}, W_{dm}, W_{dc}) =$$
$$cE'_{dm}{R'_{dm}}^{\frac{1}{2}} \cdot {\delta_{dm}}^{\frac{5}{2}} + 11\sqrt{\left(\frac{\pi^3}{32}\right)}\left(\frac{\delta_{dm}}{R_{dm}}\right)^2 \kappa_m + \pi\gamma_{dm} \cdot {\delta_{dm}}^2 - 2\pi W_{dm} R'_{dm} \cdot \delta_{dm}$$
$$+cE'_{dc}{R'_{dc}}^{\frac{1}{2}} \cdot {\delta_{dc}}^{\frac{5}{2}} + \pi\gamma_{dc} \cdot {\delta_{dc}}^2 + \eta\dot{x}6\pi R'_{dc} \cdot \delta_{dc} - 2\pi W_{dc} R'_{dc} \cdot \delta_{dc}; \quad (4.21)$$

To reduce the number of free fitting parameters, we first defined the values of the variables that were measured (or calculated) and expressed the work of adhesion of each interface $W_{ij}$, as determined by (4.10) and (4.12) respectively:

$$U_{total}(\delta_{dm}, \delta_{dc}, \gamma_{dm}) =$$
$$cE'_{dm}R'^{\frac{1}{2}}_{dm} \cdot \delta_{dm}^{\frac{5}{2}} + 11\sqrt{\left(\frac{\pi^3}{32}\right)}\left(\frac{\delta_{dm}}{R_{dm}}\right)^2 \kappa_m + \pi\gamma_{dm} \cdot \delta_{dm}{}^2 - \frac{7\pi R'_{dm}\gamma_{dc} \cdot \delta_{dm}}{4}$$
$$+cE'_{dc}R'^{\frac{1}{2}}_{dc} \cdot \delta_{dc}^{\frac{5}{2}} + \pi\gamma_{dc} \cdot \delta_{dc}{}^2 + \eta\dot{x}6\pi R'_{dc} \cdot \delta_{dc}$$
$$-\left(4\pi R'_{dc}\gamma_{dm} - \frac{9\pi R'_{dc}\gamma_{dc}}{4}\right) \cdot \delta_{dc}\,; \quad (4.22)$$

Finally, we coupled the membrane and cytosol invagination or penetration depth $\delta_{ij}$ with a simple function:

$$\delta_{dc} \to f(\delta_{dm}) = \mu \cdot \delta_{dm} + k; \quad (4.23)$$

, that reflects a mechanical coupling, where $\mu$ and k are constants that were solved for the ($\delta_{dm}$,$\delta_{dc}$) coordinates (0,0) and (7 x $10^{-8}$, 1.2 x $10^{-7}$) to 1.7 and 0 respectively. This linear relationship between $\delta_{dm}$ and $\delta_{dc}$ ensures that a critical condensate volume is conserved and that invagination values are consistent with the distribution of condensate size and membrane invagination from imaging and EM data (Idrissi, Blasco et al. 2012). This single $\delta_{ij}$ variable was henceforth referred to as δ (without any index).

We ended with a total energy function with 2 independent variables:

$$U_{total}(\delta, \gamma_{dm}) = cE'_{dm}R'^{\frac{1}{2}}_{dm} \cdot \delta^{\frac{5}{2}} + 11\sqrt{\left(\frac{\pi^3}{32}\right)}\left(\frac{\delta}{R_{dm}}\right)^2 \kappa_m + \pi\gamma_{dm} \cdot \delta^2 - \frac{7\pi R'_{dm}\gamma_{dc} \cdot \delta}{4}$$
$$+cE'_{dc}R'^{\frac{1}{2}}_{dc} \cdot (\mu\delta)^{\frac{5}{2}} + \pi\gamma_{dc} \cdot (\mu\delta)^2 + \eta\dot{x}6\pi R'_{dc} \cdot (\mu\delta)$$
$$-\left(4\pi R'_{dc}\gamma_{dm} - \frac{9\pi R'_{dc}\gamma_{dc}}{4}\right) \cdot (\mu\delta); \quad (4.24)$$

that expresses the total energy as a function of membrane invagination and interfacial tension of the endocytic condensate-membrane interface.

We explicitly defined the mechanical strain term of the model (4.1):

$$\phi \cdot \delta^{1+\varepsilon} = \begin{pmatrix} cE'_{dm}R'_{dm}{}^{\frac{1}{2}} + 11\sqrt{\left(\frac{\pi^3}{32}\right)}\left(\frac{\delta^{-\frac{1}{2}}}{R_{dm}{}^{2}}\right)\kappa_m + \pi\gamma_{dm}\cdot\delta^{-\frac{1}{2}} + cE'_{dc}R'_{dc}{}^{\frac{1}{2}}\mu^{\frac{5}{2}} \\ + \pi\gamma_{dc}\cdot(\mu\delta)^{-\frac{1}{2}} + \eta\dot{x}6\pi R'_{dc}\cdot(\mu\delta)^{-\frac{3}{2}} \end{pmatrix}\cdot\delta^{\frac{5}{2}}; \quad (4.25)$$

, where the power law variable ε of 1.5 reflects the energy penalties for parabolic shaped cytosol and membrane deformations, and the mechanical work term of (4.1):

$$\psi\cdot\delta = \left(\frac{7\pi R'_{dm}\gamma_{dc}}{4} + \mu\left(4\pi R'_{dc}\gamma_{dm} - \frac{9\pi R'_{dc}\gamma_{dc}}{4}\right)\right)\cdot\delta; \quad (4.26)$$

We minimized the function (4.24) on the $\delta_{dm}$ (1 x $10^{-9}$ m to 7 x $10^{-9}$ m) and $\gamma_{dm}$ interval (1.8 x $10^{-5}$ N•m$^{-1}$ to 7 x $10^{-5}$ N•m$^{-1}$) and obtained an energy minimum. This corresponds to an energy optimal invagination of 41 nm with a maximal $\gamma_{dm}$ of 7 x $10^{-5}$ N•m$^{-1}$. For the $\gamma_{dm}$ value of membrane invagination is favorable up to a depth of 80 nm (Figure S6). We also determined that to achieve an energy favorable membrane invagination of 70 nm (total energy $U_{total}$ equal or less than 0 J) the system requires a minimal $\gamma_{dm}$ of 6 x $10^{-5}$ N•m$^{-1}$ (Figure S6D). With this lower $\gamma_{dm}$ value, the system would reach a minimum energy with a membrane invagination of 35 nm (Figure S6D).

With the geometric data and estimates of $\gamma_{dc}$, we determined the energy required to create new interfaces $U_i$ around the condensate and the adhesive energy $U_a$ at these interfaces (Figure 6B, S6 and Methods; Eq. 4.17,4.19). At a δ value of 41 nm (corresponds to energy minimum), we summed the energy penalties (ϕ term) and estimated a total energy barrier of 2.4 × $10^{-18}$ J to deform the membrane and cytosol in contact with the endocytic condensate. This energy cost includes the elastic, viscous, and interfacial stress penalties (Figure 6B and Methods; Eq. 4.25, Table S4). The interfacial stress penalty to form the condensate/cytosol interface is 1.0 × $10^{-18}$ J and is equivalent to the membrane elastic penalties. These results are consistent with studies of artificial materials where at the 100 nm scale, surface stress can dominate elasticity in material responses to deformation (Style, Hyland et al. 2013). When we consider the energy favourable domain, our model correctly predicts the magnitude of invagination (about 40 nm to 80 nm) that is accessible for a successful invagination that leads to vesicle excision (Figure 6C and S6).

To relate the free energy on the endocytic condensate interface to density of molecular interaction on the condensate surface, we divided the adhesion energy of 3 x $10^{-18}$ J at the cytosol interface by the protein density on the condensate surface. We estimated, based on our dextran exclusion experiment, that proteins on the condensate surface are arranged in a matrix with an average mesh size of 10 nm (or less). We used an average protein filament width of 2 nm and a condensate area of 6.93 x $10^{-14}$ $m^2$ to obtain a minimum of 1.4 x $10^3$ protein segments on the condensate surface. To maximize the adhesive energy per protein exposed on the interface, we determined the minimum amount of protein on the condensate surface to be about 2 x $10^{-21}$ moles and a maximal adhesive energy density of 1.3 kJ•$mol^{-1}$. Conversely, this approach gives a maximum of 2.8 x $10^{10}$ molecules on the surface, or 4.7 x $10^{-14}$ moles of proteins, and a minimal adhesive energy of 6.3 x $10^{-5}$ J•$mol^{-1}$. To be stringent, we considered the former.

#### Statistical analysis and software for microscopy

Software used in microscopy measurements and microscopy data analysis are also outlined in the methods details. For dSTORM images in Figure 1B, center of mass for each event was calculated and we reconstructed images in Wave Tracer before further analysis in Metamorph (Molecular Devices). For the measurements of fluorescent probe uptake (Figure 4A,C,D and S1E-F) and cell sizes under different osmotic pressures (Figure S1A), image analysis and signal automated segmentation was performed with the InCell Developer software (GE healthcare life sciences) and the data was further analyzed and plotted in the R environment. Quantification of FM4-64 membrane probe (Figure 2C and 6C) was performed with Metamorph image analysis software (Molecular Devices). When cell populations in Figure 2C and Figure 3D were compared for quantified fluorescence, we applied a Welch's two-sided t-test with sample sizes (n = 100) to achieve a power greater than 0.9 with a 95% confidence. Statistical analysis was done with the R software package. For particle tracking and mean squared displacement analyses in Figure S4, we used the Wave Tracer plugin with the Metamorph image analysis software (Molecular Devices) to localize fluorescent foci centroid position through a wavelet algorithm and tracks particles in times stacks to calculate particles movement. Kymographs in Figure S1 were also generated with the Metamorph image analysis software (Molecular Devices). Analysis of the images intensity (I) fluctuations and segmentation of regions of interest (ROI) for fluorescence recovery after photobleaching (FRAP) experiments (Figure 2E, 5E-F and S5C-D) were performed on the LASX imaging software (Leica). FRAP data was plotted using subroutines of the R package. Optical tweezers measurements were performed with in house Labview (National Instruments) routines and the data was analyzed with a custom Matlab (MathWorks) code.

#### Theoretical model based on elastic and adhesive contact mechanics

Calculations and graphics of the elastic, viscous, surface and adhesive energies in our model were performed in either Maple or the R software environment. The parameters, variables and relationships we used in the calculation are listed in Table S2 and S3. A summary of the invagination and energy results we obtained with these specific parameters are in Table S4.

### Supplementary Figures

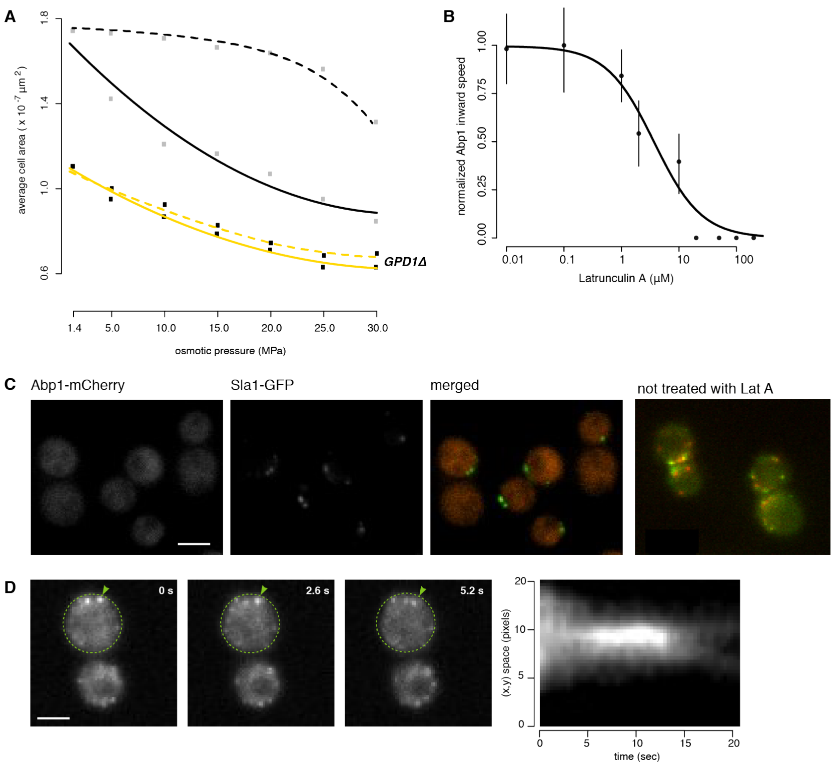


**Fig. S1 *GPD1Δ* cells undergo normal endocytosis in the absence of actin polymerization (Latrunculin A-treated).**

(A) To detect turgor pressure and cell size adaptation to osmotic shock in wild type (black) and *GPD1Δ* strain (yellow), we monitored by fluorescent microscopy the cross-sectional area (µm$^2$) of shocked cells (solid lines) and adapted cells (dashed lines) in water-glycerol binary solutions from 1.4 MPa to 30 MPa. Points represent mean area values (n=200 cells).

(B) Dose-response of normalized Apb1-mCherry inward speed as a function of Latrunculin A (Lat A) concentration (mean ± sd; n = 12; logistic fit). Apb1-mCherry tracks in time were analyzed from kymographs of the Lat A-treated cells. Note that Abp1-mCherry structures disappear at 20 µM Lat A and above.

(C) F-actin polymerization is disrupted by Lat A. Assessment of Abp1-mCherry actin structures by fluorescence microscopy in cells treated with 20 µM Lat A for 5 min. Maximal intensity projections of Z-stacks of cells after and without treatment are shown, Sla1-GFP (green signal) and Abp1-mCherry (red signal). Scale bar, 2 µm.

(D) Time progression of Sla1-YFP-labeled puncta (arrow) in *GPD1Δ* cells (dashed contour) treated with 20 µM Lat A (effective concentration determined in B-C). Elapsed time relative to puncta appearance is indicated is top right corners. Scale bar, 2 µm. The kymograph in the right panel shows spatiotemporal progression of the Sla1-YFP-labelled puncta identified by the arrow.

**A**

| | Control | | 0.5% HD | | 1% HD | | 1.5% HD | | 2.0% HD | | 2.0% HT | |
|---|---|---|---|---|---|---|---|---|---|---|---|---|
| | before | after | before | after | before | after | before | after | before | after | before | after |
| Trial 1 | 31 | 28 | 32 | 31 | 32 | 29 | 33 | 31 | 30 | 27 | 30 | 28 |
| Trial 2 | 25 | 23 | 33 | 30 | 27 | 26 | 33 | 33 | 26 | 20 | 26 | 23 |
| Trial 3 | 35 | 33 | 31 | 32 | 36 | 33 | 31 | 30 | 33 | 31 | 32 | 27 |
| Trial 4 | 27 | 25 | 29 | 28 | 24 | 23 | 32 | 28 | 28 | 26 | 26 | 25 |
| Mean±S.D. | 29±4 | 27±4 | 31±2 | 30±2 | 30±5 | 28±4 | 32±1 | 30±2 | 29±3 | 26±5 | 28±3 | 26±2 |

**B**

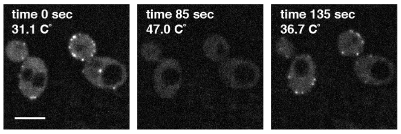


**Fig. S2 Sla1-labelled puncta are reversibly thermo-sensitive.**

(A) Cell viability is unaffected by heating and cooling cycles and HD treatment.

(B) Single cell assay confirms that Sla1-labelled puncta returned throughout the temperature cycles. Scale bar, 4 μm.

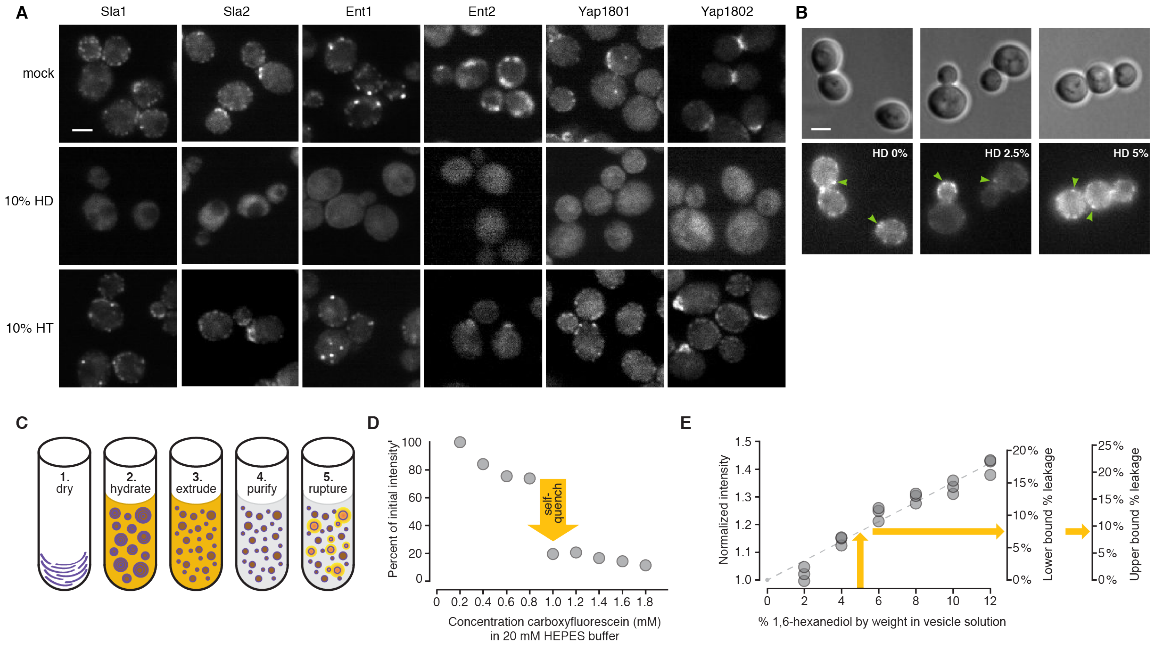


**Fig. S3 PLD-containing proteins form puncta are reversibly sensitive to 1,6-hexanediol (HD).**

(A) Fluorescence images of GFP-tagged Sla1, Sla2, Ent1, Ent2, Yap1801 and Yap1082 puncta 5 min after treatment with either DMSO, 1,6-hexanediol or 1,2,3-hexanetriol, images where acquired with InCell6000 confocal microscope. Scale bar 4 µm.

(B) Brightfield (top) and fluorescence (bottom) images of *GPD1Δ* cells Syp1-mCherry cells treated with 05, 2.5 and 5% HD, show that Syp1-mCherry foci are not dissolved by HD (arrows). Scale bar 4 µm.

(C) Protocol for determining vesicle leakage by 1,6-hexanediol. Vesicles containing carboxyfluorescein were produced through drying (Step 1), hydration (Step 2), and extrusion (Step 3). Dried lipids were hydrated with a 20 mM carboxyfluorescein solution (~20x the self-quenching concentration in panel B). Unencapsulated carboxyfluorescein was removed on a size exclusion column (Step 4). Vesicle rupture or leakage causes carboxyfluorescein concentrations to fall below the self-quenching limit, resulting in increased fluorescence (Step 5).

(D) Carboxyfluorescein self-quenches at concentrations ≥ 1.0 mM in solution as seen by the sharp decrease in fluorescence intensity at 1.0 mM. Intensity of the first data point was set to 100 (in arbitrary units).

(E) Vesicle leakage due to 1,6-hexanediol is minor. Carboxyfluorescein fluorescence intensity of vesicle solutions treated with 1,6-hexanediol (2 to 12% by weight). Standard deviations were smaller than the symbols in the graph (n = 3; repeated 3 times). Data were normalized such that a solution with no 1,6-hexanediol has an intensity of 1.0. A linear fit constrained to pass through (0, 1.0) is shown by the dashed line. Consistent with the data in panel B, full de-quenching resulted in a 2.95- to 3.34-fold increase in fluorescence. Propagation of this range of multipliers leads to the conclusion that for vesicles in 5% 1,6-hexanediol solutions (vertical arrow) only 7.5% (horizontal arrow, lower bound) to 9.0% (upper bound) of carboxyfluorescein leaks out of vesicles.

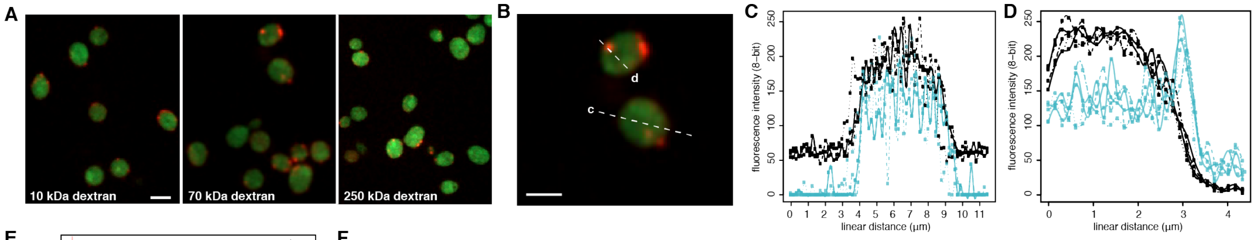


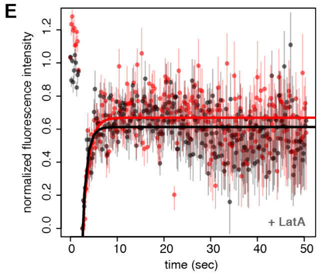


F

| dextran-FITC | LatA / HD | 10 kDa | 70 kDa | 250 kDa |
|---|---|---|---|---|
| size | | 2.4 nm | 5.8 nm | 10.4 nm |
| mobile fraction in droplet | + / - | 0.71 ± 0.01 | 0.77 ± 0.03 | 0.35 ± 0.07 |
| mobile fraction in cytosol | + / - | 0.71 ± 0.01 | 0.94 ± 0.04 | 0.89 ± 0.03 |
| mobile fraction in droplet | + / + | nd | 1.23 ± 0.08 | 0.80± 0.13 |
| mobile fraction in cytosol | + / + | nd | 1.27 ± 0.11 | 0.86± 0.08 |

**Fig. S4 Colocalization of different sized dextran-FITC revealed that endocytic puncta exclude objects of diameters larger than 10.4 nm**

(A) 2-color fluorescent images of osmoporated dextran-FITC of different sizes in Sla1-mCherry treated with Lat A, Scale bar is 4 µm.

(B) 2-color fluorescent images of dextran-FITC of 5.8 nm in size osmoporated into *GPD1Δ* Sla1-mCherry cells treated with Lat A, Scale bar is 4 µm. Examples of line scans (dashed) that were performed either outside (c) or across (d) Sla1-mCherry foci.

(C) Multiple line scans of 5.8 nm dextran-FITC fluorescent signal (black) outside Sla1-mCherry cortical regions (blue). Fluorescence intensity patterns thus show the fluorescence of a line across the cytoplasm that does not include any concentrated foci of Sla1-mCherry.

(D) Multiple line scans of 5.8 nm dextran-FITC fluorescent signal (black) within Sla1-mCherry cortical regions (blue) shows partial exclusion of these molecules.

(E) Fluorescence recovery after photobleaching (FRAP) of the bleached 2.4 nm dextran-FITC within a Sla1-mCherry punctum (red) and neighbouring cytosol regions (black) without Sla1 signal. *GPD1Δ* cells Sla1-mCherry were treated with 20 µM Lat A. Data points (mean ± SEM; n = 10 cells) were fitted to a single term recovery function (Methods).

(F) Summary of mobile fractions determined for 2.4 nm, 5.8 nm and 10.4 nm dextran-FITC in either the endocytic puncta region or neighbour cytosol (mean ± se estimated from fits).

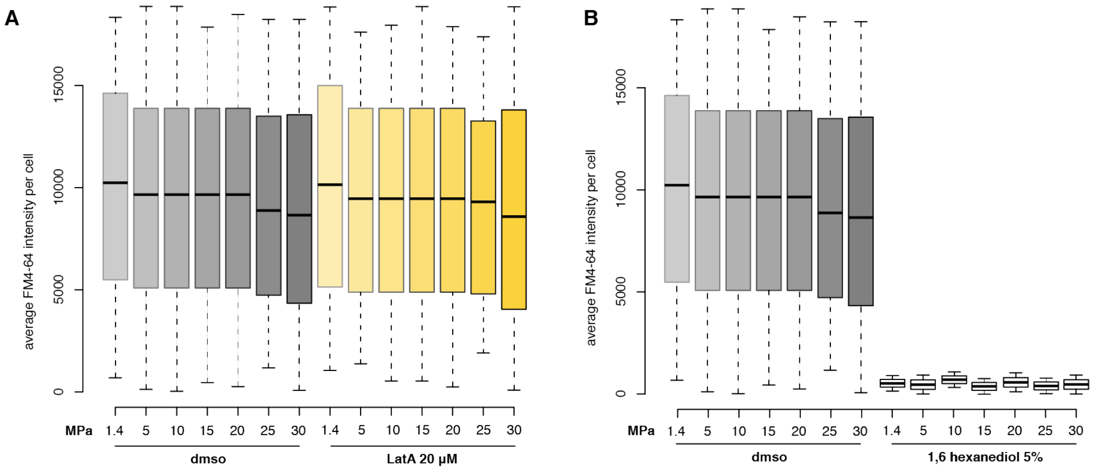


**Fig. S5. Sla1-labelled puncta are characterized by thermal and solute responsive non-covalent interactions and that these puncta are essential for actin-independent endocytosis in *GPD1Δ* cells**

(A) Membrane-associated FM4-64 fluorescent probe uptake was quantitatively assessed by fluorescence imaging (box center line, median; box limits, upper and lower quartiles; whiskers, 1.5x IQR; crosses, outliers). *GPD1Δ* strains undergo normal endocytosis when treated with Lat A, (B) but not when treated with 1,6-hexanediol, n = 900 cells for each condition.

A

```
SLA1 PLD      -------------------------------------------------------------   0
SUP35 NM(Q)   MSDSQQGQ--QQQQYQQYSQQGQQQQGQQRYQGYQAYQAQAQPAGGYYQQYQGYSGYQQG   58
ENT1 PLD      MLQRM--------------QQQ---QGQQQ-----------LQQPMYYDIFGNP--ITPE   30
ENT2 PLD      MLQELQRLQKQQQSLSQFQAPL---QQQQP-----------QQQPAYYDIFGNP--ISQD   44

SLA1 PLD      ----------------------------------------MCGGAQFPQT--S----   11
SUP35 NM(Q)   GYQQYQPDA-----GYQQQYQPQG-GYQQYQPQGGYQQQFQ-PQGGRGQYKQFQYQQQ--   109
ENT1 PLD      EYAQFQLQQQQQQQQQ------------LQQQPMYYDVFGNP-ITPEELAQFQQQQQLQ   77
ENT2 PLD      EYLQYQYQQDQEQAMAQQRWLDQQQEQQQLAEQ-QYFQQQQQAAAAASALQQQQTAANMQ   103
                                                                   *

SLA1 PLD      ----FNVQGQQQLPTGSILPVQKTANGLIS-----------------ANTGVSMPTVQRT   50
SUP35 NM(Q)   -LQGYQAGFQPQSQGMSLQDFQKQQKQAAPKPKKTLKLV--------SSSGIKLANATKK   160
ENT1 PLD      -EQQYLASMQQQQQAMSNNPFAKSEQSSSS------------------------------   106
ENT2 PLD      QQQQQPADFQQPLPTGSNNPFSMDNLERQKQEQQHAQLQRQQEEARQQQEQLKLQQLQRQ   163
                    .  *      *   .

SLA1 PLD      GGTMIPQTSFGVSQQLTGGAMMTQPQNTGSAMMPQT--SFNA-VPQ-----------ITG   96
SUP35 NM(Q)   VGTKPAE-----SDKKEEEKSA----ETK---EP-TKEPTKVEEPVKKEEKPVQTEEKTE   207
ENT1 PLD      -------------PKRN---QL------VAASSPQQLQQQKQQEPLIQN--------RTG   136
ENT2 PLD      QQEEAQL-----HQKRQEEAQL----QQQQAQLLQQQAQFQQQQPLKQT--------RTG   206
                          :                         :   *             *

SLA1 PLD      -GAMMPQTSFNALPQVTGGAMMPLQRTGGALNTF-NTGGAMIPQTSFSSQAQNTGGFRPQ   154
SUP35 NM(Q)   EKSELPKV----------EDLKISEST-------HNTNNANV----TS-----ADALIKE   241
ENT1 PLD      NQSMTDKY----------SKLNELLATGTGIDTFGNVGEARIPAQHTK-----TGTFINS   181
ENT2 PLD      NQSISDKY----------SDLNTLLATGTGIDTFGNTGEARIPAQHTK-----TGTFINS   251
                :   :           :     *        *.. * :      .     :. :  .

SLA1 PLD      SQFGLTLQKTGGIAPLNQNQFTGGAMNTLSTGGVLQQQQPQTMNTFNTGGVMQELQMMTT   214
SUP35 NM(Q)   QEEE--------------------------------------------------------   245
ENT1 PLD      QGTGY-------------------------------------------------------   186
ENT2 PLD      QGTGY-------------------------------------------------------   256
              .

SLA1 PLD      FNTGGAMQQPQMMNTFNTDGIMQQPQMMNTFNTGGAMQQPQQQALQNQPTGFGFGNGPQQ   274
SUP35 NM(Q)   ---------------VDDEVVNDN------------------------------------   254
ENT1 PLD      ---------RQ-----VSDDPNHNPFLNSQY-----TG-LPSTSVVPTQTGYGFGNQSQQ   226
ENT2 PLD      ---------KQ-----VTNEPKNNPFLSNQY-----TG-LPSTNIVPTQTGYGFGNQPQS   296
                               :    ::

SLA1 PLD      SRQANIFNATASNPFGFD------------------------------------------   292
SUP35 NM(Q)   ------------------------------------------------------------   254
ENT1 PLD      QSQNN----GSNNR-GYTLIDL--------------------------------------   243
ENT2 PLD      PPTNS----PQQNPTGISYSQPQQQQQPQQQPQYMQNFQQQQPQYAQNFQQQPQYTQNYQ   352

SLA1 PLD      ------------------------------------   292
SUP35 NM(Q)   ------------------------------------   254
ENT1 PLD      ------------------------------------   243
ENT2 PLD      QQPQYIQPHQQQQQQQQQQQQQQGYTPDQGVSLIDL   388
```

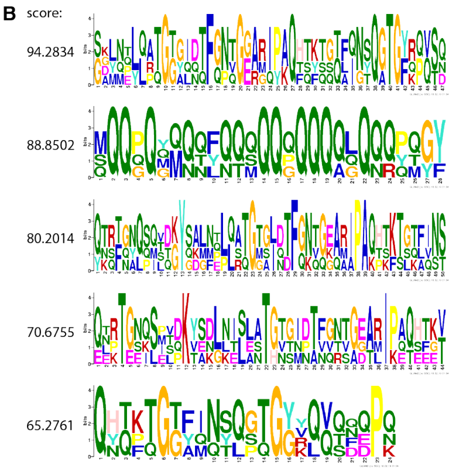


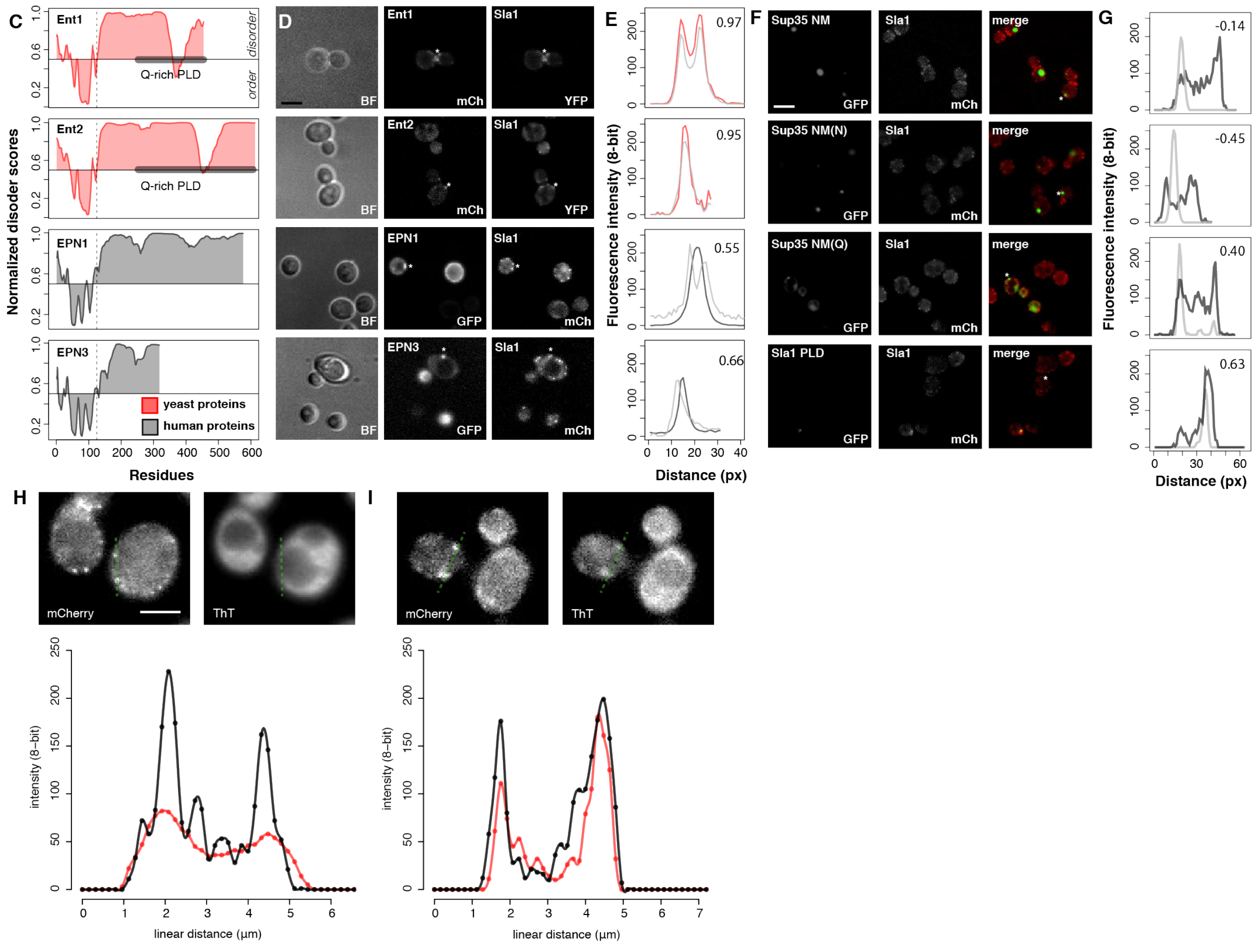

**Fig. S6 Sequence and colocalization analysis of PLDs from human Epsin orthologs and SUP35 NM sequence variants.**

(A) Sequence alignment of Ent1 PLD, Ent2 PLD, Sla1 PLD and Sup35 NM(Q) with Clustal Omega(18).

(B) Search for common binding motifs in the PLD sequences of Ent1, Ent2 and Sla1 and the sequence of Sup35 NM(Q) using motif discovery tool GLAM2(19).

(C) PNDR-fit score (VLS2) for yeast Ent1 or Ent2 (red) and human EPN1 or EPN3 (black) proteins show conservation of the disorder along the sequence of these yeast (red) and human (grey) homologs.

(D) Imaging of Ent1, Ent2, EPN1 and EPN3 proteins with GFP fluorescent tags and assessment of their colocalization with Sla1-labeled endocytic puncta (mCherry tag). Scale bar, 5 μm.

(E) Line scans (performed as indicated with star in B) confirm, that neither EPN1 nor EPN3 colocalize with Sla1 foci (grey lines) in yeast cells. Pearson correlation values are given to confirm if the signals colocalize.

(F) Imaging of Sup35 NM variants and Sla1 PLD proteins with GFP fluorescent tags and assessment of their colocalization with Sla1-labeled endocytic puncta (mCherry tag). Scale bar, 5 μm.

(G) Line scans (performed as indicated with star in C) confirm, that neither Sup35 NM, Sup35 NM(N) with all Qs mutated to Ns nor Sup35 NM(Q) with all Ns mutated to Qs (black lines) colocalize with Sla1 foci (grey lines) in yeast cells, unlike Sla1 PLD (black) that colocalize with endogenous Sla1 proteins (grey). Pearson correlation values are given to confirm if the signals colocalize.

(H) We observed no colocalization of Thioflavin T (ThT) with Sla1-mCherry puncta. Fluorescence microscopy of Sla1-mCherry (upper left), ThT stain (upper right) and line scan analysis (lower panel) for the dashed line in green. Scale bar, 2 μm.

(I) Fluorescence microscopy of Sup35-mCherry (upper left), ThT stain (upper right) and line scan analysis (lower panel) for the dashed line in green. Scale bar, 2 μm.

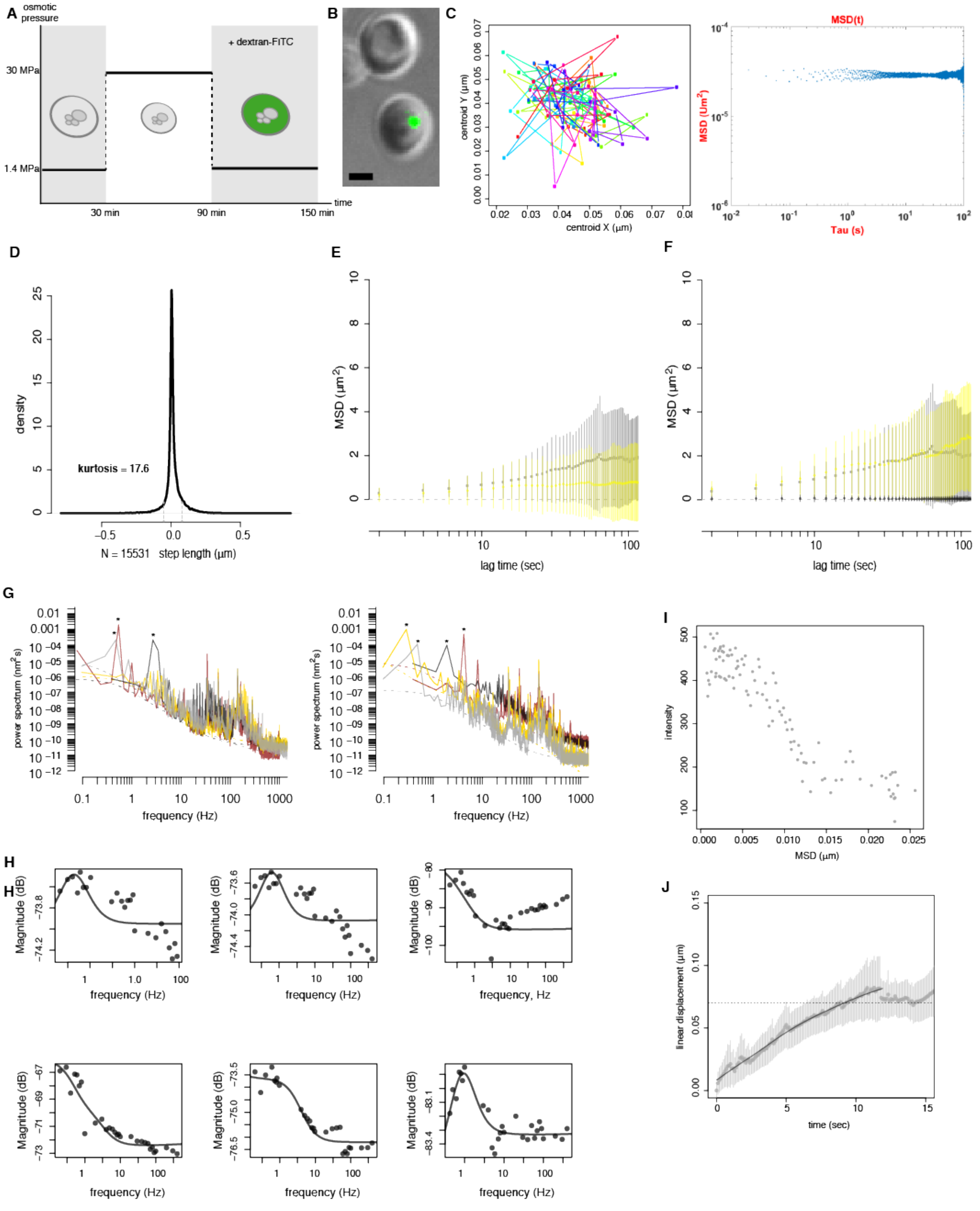


**Fig. S7 Dynamic mechanical analysis of cytosol with optical tweezers.**

(A) Schematic representation of the two-cycle osmotic shock used to osmoporate 200 nm polystyrene beads into haploid yeast *GPD1Δ* cells treated with Lat A. The osmoporation treatment induces a senescent-like state in most cells, but we could rescue a small fraction of cells that continue to divide in rich YPD medium.

(B) Images of Sla2-Dronpa3 (green foci) cells with osmoporated beads (left panel). Scale bar, 2 μm.

(C) Passive 2D displacement in x and y of 200 nm polystyrene beads determined by centroid tracking (left panel) and mean square displacement (right panel). The polystyrene beads we incorporated into cells were confined within the cytosol with a MSD close to that of technical noise.

(D) Distribution of step length of expressed microNS-GFP particles determined by centroid tracking in untreated cells.

(E-F) MSD of microNS particles in untreated (grey) and osmoporated (yellow) cells as a function of lag time (seconds). Data points represent mean ± standard deviation of n = 300 traces in untreated *versus* osmoporated cells, panel E shows raw data and panel F shows data filtered for caged particles with MSD < 1 represented in black.

(G) Viscoelastic properties of the cytosol of Lat A treated *GPD1Δ* cells measured using the response of 200 nm polystyrene beads to sinusoidal oscillations of the tweezers and the high frequency domain of the power spectrum associated with thermal motions of the bead. Power spectra of samples are coded with distinct colors. Below frequencies of 500 Hz, the power spectra show fluctuations from tweezer oscillations (marked *), cellular processes and sample vibration.

(H) Magnitude of the response (dB) of the bead displacement to sinusoidal oscillations of the optical trap position. Lines indicate global fits of the power spectra and response magnitudes to a model that describes the viscoelasticity of a crosslinked polymer network (Methods).

(I) Centroid tracking within a confocal volume of Sla1-YFP foci in *GPD1Δ* mutant strains treated with Lat A. Single Sla1-YFP puncta fluorescence intensity as a function of mean square displacement in a confocal volume as determined by spinning disk pinhole size. Images were acquired at 20 frames per second. Only foci with a decreasing fluorescence bleaching were selected to measure their displacement.

(J) Linear displacement of Sla1 foci within a confocal volume as a function of time. Mean ± standard deviation of displacement is shown for n > 50 per time point. Total traces n = 275; note that traces do not have the same length. The average displacement is about 7.4 nm per second from polynomial fit (black curve). Horizontal dashed line indicates an invagination of 70 nm, minimal depth required for the constriction and scission steps.

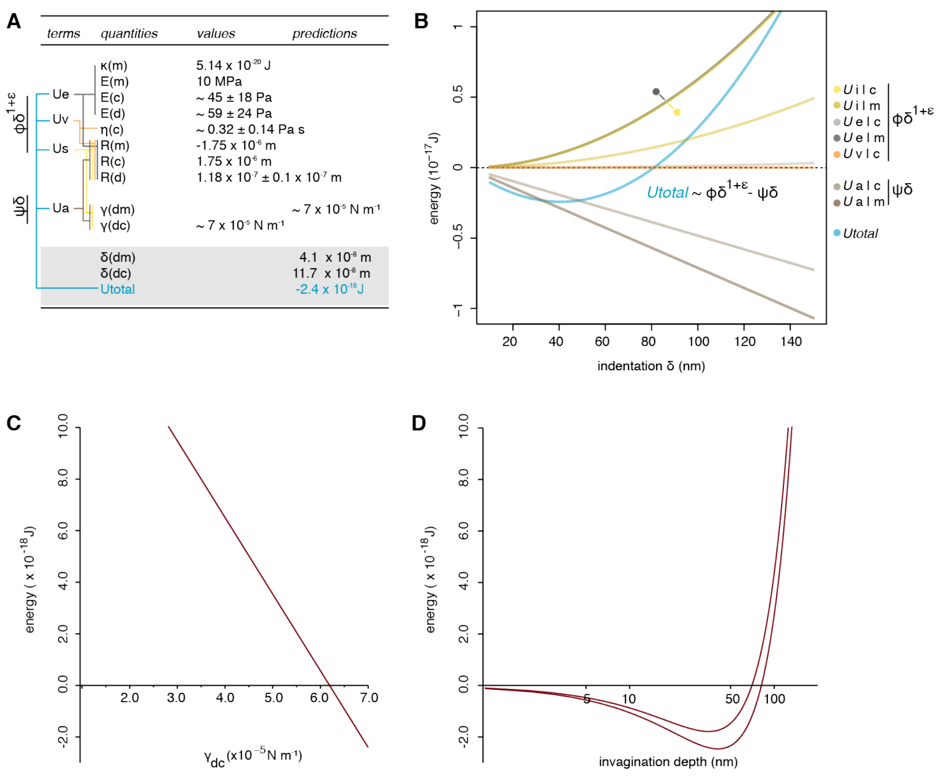


**Fig. S8 Endocytic condensates apply mechanical stress that deform the membrane.**

(A) Values of bending, elastic, viscosity, surface tension and geometry quantities used in the calculation are recapitulated. Colour coded lines indicate which quantities were used to calculate the respective energies $U_x$, that in turn compose the ϕ and ψ terms of Equation 1.

(B) Equation (1) (insert) was used to calculate the energy penalties and contributions at the cytosol and membrane interfaces with the endocytic condensate. Total energy of the system (blue), energy penalties (yellow, orange and grey) and energy contributions (brown) are presented as a function of membrane invagination (δ). Colour legend (right) specifies energy traces $U_{xly}$, where x= (E), elasticity or (v), viscosity of either the cytosol (c) or membrane (*m*) and (i) is interfacial tension and (a) is adhesion of cytosol or membrane with the endocytic condensate. Quantities used to calculate energies are detailed in Table S3 and S4.

(C) Plot of total energy (y axis) as a function of condensate-cytosol surface tension (x axis) shows that the energy favorable values of $\gamma_{dc}$ range between 6.2 x$10^{-5}$ N•$m^{-1}$ to 7 x$10^{-5}$ N•$m^{-1}$.

(D) 2D representation of the energy and depth accessible to a successful invagination. With the minimal (top curve) and maximal (lower curve) values of $\gamma_{dm}$ our model predicts precise range of favorable δ (x axis) that minimizes total energy (y axis).

### TABLES

**Table S1.** Strains and primers used in this study.

| *name* | *genotype* | *source* |
|---|---|---|
| BY4741 | MATa his3Δ1 leu2Δ0 met15Δ0 ura3Δ0 | |
| GPD1Δ | BY4741 gpd1Δ::KanMX | YKO |
| Ent1-GFP | B4741 ent1-GFP::His3MX | GFP |
| Ent2-GFP | B4741 ent2-GFP::His3MX | GFP |
| Pan1-GFP | B4741 pan1-GFP::His3MX | GFP |
| Sla1-GFP | B4741 sla1-GFP::His3MX | GFP |
| Sla2-GFP | B4741 sla2-GFP::His3MX | GFP |
| Yap1801-GFP | B4741 yap1801-GFP::His3MX | GFP |
| Yap1802-GFP | B4741 yap1802-GFP::His3MX | GFP |
| Ent1-PLDΔ | BY4741 ent1-PLDΔ-Venus::HygMX | this study |
| Ent1-venus | B4741 ent1-Venus::HygMX | this study |
| Ent2-PLDΔ | BY4741 ent2-PLDΔ-Venus::HygMX | this study |
| Ent2-venus | B4741 ent2-Venus::HygMX | this study |
| ENT2Δ | BY4741 ent2Δ::KanMX | YKO |
| GPD1Δ Sla1- mCherry | GPD1Δ sla1-mCherry::HygMX | this study |
| GPD1Δ Sla1-venus | GPD1Δ sla1-venus::HygMX | this study |
| SLA1Δ | BY4741 sla1Δ::KanMX | YKO |
| GPD1Δ Syp1- mCherry | GPD1Δ syp1-mCherry::HygMX | this study |
| Sla1-GFP Abp1- mCherry | BY4741 sla1-GFP::His3MX abp1-mCherry::HygMX | this study |
| Sla1-mCherry | BY4741 sla1-mCherry::HygMX | this study |
| Sla1-PLDΔ | BY4741 sla1-PLDΔ-Venus::HygMX | this study |
| Sla1-venus | B4741 sla1-Venus::HygMX | this study |
| Sla2-GFP Sla1-mCherry | B4741 sla2-GFP::His3MX sla1-mCherry::HygMX | this study |
| Sup35-mCherry | BY4741 sup35-mCherry::HygMX | this study |
| Yap1801-PLDΔ | BY4741 yap1801f-PLDΔ-Venus::HygMX | this study |
| Yap1801-venus | B4741 yap1801-Venus::HygMX | this study |
| Yap1802-PLDΔ | BY4741 yap1802-PLDΔ-Venus::HygMX | this study |
| Yap1802-venus | B4741 yap1802-Venus::HygMX | this study |

**Table S2.** Parameters and variables used in our model.

| *parameters* | *definition* | *approximative value* | *note* | *source* |
|---|---|---|---|---|
| $E_m$ | membrane elastic modulus | 1 x10$^{7}$ Pa | *estimate* | Landau 1986 |
| $\kappa_m$ | membrane bending modulus | 12.5•$K_BT$ | | Harmandaris 2006 |
| $v_m$ | membrane poisson's ratio | 0.45 | *from 0.1 to 0.5* | Zhang 2013 |
| $\delta_m$ | membrane indentation | 5 x10$^{-8}$ m | from 2.5 x 10$^{-8}$ to 5 x 10$^{-8}$ m | EM |
| $R_m$ | membrane radius | -1.75 x 10$^{-6}$ m | *negative curvature* | *this study* |
| $E_c$ | cytosol elastic modulus | 45 Pa | ± 18 Pa (value at 1 Hz) | *this study* |
| η | cytosol viscosity | 0.35 Pa•s$^{-1}$ | ± 0.14 Pa•s$^{-1}$ *(value at 0.5 Hz)* | *this study* |
| $v_c$ | cytoplasm poisson's ratio | 0.45 | *from 0.1 to 0.5* | Zhang 2013 |
| $\delta_c$ | indentation cytosol | 1.18 x 10$^{-7}$ m | ±6 x 10$^{-9}$m | *this study* |
| $R_c$ | membrane radius | 1.75 x 10$^{-6}$ m | *positive curvature* | *this study* |
| $E_{drop}$ | droplet elastic modulus | 59 Pa | ± 24 Pa (value at 1 Hz) | *this study* |
| $2a_{drop}$ | droplet contact diameter | 2.09 x 10$^{-7}$ m | ±1 x 10$^{-8}$m | *this study* |
| $\delta_{drop}$ | indentation droplet | 1.18 x 10$^{-7}$ m | ±10 x 10$^{-9}$m | *this study* |
| $R_{drop}$ | droplet radius | 9.3 x 10$^{-7}$ m | ~$a^2/R$ | *Hertz* |
| θ | droplet contact angle | 96.7° | $\theta = 2\ \arctan(\delta_{drop}/a_{drop})$ | Young |

* EM refers to electron microscopy data (20) and Hertz refers to Hertz contact theory (21). Landau 1986 refers to estimation of membrane elastic modulus from the bending modulus and Poisson's ratio (22), Harmandaris 2006 (9) and Zhang 2013 (11).

**Table S3.** Constants and equations used in the elasto-adhesion model for the deformation of the membrane by protein droplets on cortical sites.

| *variables* | *definition* | *approximative value* | *note* | **source** |
|---|---|---|---|---|
| *mechanical stress method* | | | | |
| $\varepsilon$ | mechanical deformation | *nd* | $\delta_x/R$ | Hooke |
| $\varepsilon^{\cdot}$ | cytosol deformation rate | 0.004 $s^{-1}$ | | *this study* |
| $E_{cell}$ | cell apparent elastic modulus | 1000 Pa | | AFM |
| $\sigma$ | mechanical stress | 1181Pa | $\sigma = \varepsilon E + \varepsilon\eta$ | Kelvin-Voigt |
| $\Delta P$ | pressure difference | 1181Pa | $\Delta P = \sigma$ | Laplace |
| H | interface mean curvature | 8.5 x$10^{6}$ $m^{-1}$ | 1/R | Laplace |
| $\gamma_{dc}$ | droplet-cytoplasm surface tension | 7 x$10^{-5}$N•$m^{-1}$ | $\gamma = \Delta P/(2H)$ | Young-Laplace |
| *elasto-adhesive contact method* | | | | |
| c | constant | 0.92 | $8/(5\sqrt{3})$ | JKR |
| $E^{*}_{dc}$ | cyto. vinterface equivalent elastic modulus | 32 Pa | $1/E^{*}_{c} = (1-v_c^2)/E_c + (1-v_m^2)/E_m$ | Hertz |
| $E^{*}_{dm}$ | membrane interface equivalent elastic modulus | 75 Pa | | Hertz |
| $R_{dm}$ | equivalent radius | 1.27 x$10^{-7}$ m | $1/R = 1/R_m + 1/R_d$ | Hertz |
| $R_{dc}$ | equivalent radius | 1.11 x$10^{-7}$ m | $1/R = 1/R_c + 1/R_d$ | Hertz |
| $\delta_c$ | cytoplasm indentation | *see table S3* | $f(\delta_m) = \mu + \omega\delta_m$ | *this study* |
| $\mu$ | constant | 1.65 | | *this study* |
| k | constant | 0 | | *this study* |
| $W_{dm}$ | droplet-membrane work of adhesion | 6.15 x$10^{-5}$ N•$m^{-1}$ | $W_{dm} = \gamma_{dc}(1+\cos\theta)$ | Young Dupré |
| $W_{dc}$ | droplet-cytosol work of adhesion | 6.15 x$10^{-5}$ N•$m^{-1}$ | $= \gamma_{cm} + \gamma_{dm} - \gamma_{dc}$ predicted from model *(1 x $10^{-5}$ to 7 x$10^{-5}$N•$m^{-1}$)* | *this study* |
| $\gamma_{dm}$ | droplet-membrane surface tension | 6.15 x$10^{-5}$ N•$m^{-1}$ | $\gamma_{dm} < \gamma_{dc}$ *(hydrophobic)* $\gamma_{dm} \sim (W_{dc} + \gamma_{dc})/2$ | Young-Dupré |
| $\gamma_{cm}$ | cytosol-membrane surface tension | 5.75 x$10^{-5}$ N•$m^{-1}$ | $\gamma_{cm} = \gamma_{dm} + (\gamma_{dc}\cos\theta)$ | Young |

* not determined (nd). AFM refers to atomic force microscopy data (12).

**Table S4.** Summary of the indentations and energies predicted with our elasto-adhesive contact model.

| *variable* | *definition* | *approximative value* | *in kT* | *source* |
|---|---|---|---|---|
| $\delta_m$ | membrane invagination | 4.06 x$10^{-8}$ m | *nd* | *this study* |
| $\delta_c$ | cytoplasm deformation | 1.17 x$10^{-7}$ m | *nd* | *this study* |
| $U_{total}$ | total energy | -2.4 x$10^{-18}$ J | *-590•kT* | *this study* |
| $U_{penal}$ | total energy penalties | 2.4 x$10^{-18}$ J | *590•kT* | *this study* |
| $U_{em}$ | corrected elastic energy at droplet-membrane interface | 1 x$10^{-18}$ J | 250•*kT* | *this study* |
| $U_{ec}$ | elastic energy at droplet-cytoplasm interface | 1.2 x$10^{-20}$ J | *3•kT* | *this study* |
| $U_{\gamma m}$ | surface energy at droplet-membrane interface | 3.6 x$10^{-19}$ J | 88•*kT* | *this study* |
| $U_{\gamma c}$ | surface energy at droplet-cytoplasm interface | 1 x$10^{-18}$ J | *249•kT* | *this study* |
| $U_{vc}$ | viscous friction energy | 2.5 x$10^{-21}$ J | 1•kT | *this study* |
| $U_{adh}$ | total adhesion energy | 4.9 x$10^{-18}$ J | 1180•*kT* | *this study* |
| $U_{am}$ | adhesion energy at droplet-membrane interface | 2 x$10^{-18}$ J | 477•*kT* | *this study* |
| $U_{ac}$ | adhesion energy at droplet-cytoplasm interface | 2.9 x$10^{18}$ J | 703•*kT* | *this study* |

**Movie S1**. Pulse-chase experiments with HD showed that HD-dependent dissolution of Sla1 puncta was reversible. 2-color time-lapse fluorescence images of Sla1-GFP Abp1-mCherry cells treated with HD-containing media at 10 min time point, and then with fresh media without HD at 15 min time point.
